# Azacitidine Response in Myelodysplastic Syndromes is Marked by NK-like CD8 T-Cell Expansion and CXCL12+ Reticular Cell Remodeling

**DOI:** 10.64898/2026.07.07.736922

**Authors:** Henry R. Hampton, Anne Pan, Michael Carnell, Bofei Wang, Diana Shinko, Maria Kasherman, Iveta Slapetova, Swapna Joshi, Mary N.T. Nguyen, Feng Yan, Sarah Davidson, Nick F.Y. Choi, Jason W.H. Wong, Nicodemus Tedla, Devendra K. Hiwase, Magnus Tobiasson, Mark N. Polizzotto, Helen M. McGuire, Hussein A. Abbas, Asif Javed, Jake Olivier, Julie A.I. Thoms, Christopher J. Jolly, John E. Pimanda

## Abstract

Myelodysplastic syndromes (MDS) are driven by somatic mutations in hematopoietic stem and progenitor cells (HSPCs), leading to clonal expansion and ineffective hematopoiesis. Hypomethylating agents (HMAs; azacitidine or decitabine) are the standard of care for higher-risk MDS. However, their effects on the bone marrow (BM) microenvironment, and the extent to which these changes correlate with clinical response, remain poorly understood. We performed longitudinal analyses of BM aspirates, trephine biopsies, and peripheral blood samples from MDS patients treated with azacitidine in a clinical trial (NCT03493646), integrating CyTOF, 5′ single-cell RNA and TCR sequencing, plasma proteomics, and multiplex immunofluorescence microscopy to characterize changes associated with azacitidine response. Clinical responders showed expansion of GzmB⁺CD56⁺CD8⁺ T cells together with increased type I and type II interferon signaling within the T-cell compartment. Responders also exhibited marked alterations in circulating platelet– and myeloid-derived factors with the potential to remodel the BM niche. Spatial analyses revealed expansion of “neighborhoods” enriched for CXCL12-abundant reticular cells and CD8 T cells in responders, whereas HSPC-enriched neighborhoods were largely unchanged. In contrast, several HSPC-enriched neighborhoods expanded in non-responders. These microenvironmental changes were accompanied by evidence of enhanced myelopoiesis in clinical responders. Our findings support a model in which azacitidine response extends beyond direct effects on malignant hematopoietic cells to involve coordinated remodeling of the BM microenvironment which may be reinforced by platelet– and myeloid-derived signals that establish a feed-forward circuit promoting productive hematopoiesis.

**One Sentence Summary:** Longitudinal bone marrow profiling reveals stromal and immune correlates of azacitidine response in myelodysplastic syndrome.

## INTRODUCTION

Myelodysplastic syndromes (MDS), also known as myelodysplastic neoplasms, arise from accumulation of somatic mutations in hematopoietic stem cells (HSCs), leading to clonal expansion and impaired differentiation, multilineage cytopenias, and increased risk of progression to acute myeloid leukemia (AML) (*1*). The median age of presentation is approximately 75 years (*2*), and as a result, most patients are ineligible for allogeneic stem cell transplantation, the only potentially curative treatment. Hypomethylating agents (HMAs) such as 5-azacitidine (AZA) and 5-aza-2′-deoxycytidine (dAZA; decitabine) remain the standard of care for transplant-ineligible patients. While HMAs can prolong survival and delay progression to AML, responses are variable and rarely durable (*3, 4*). Furthermore, the mechanisms by which HMA treatment leads to improved hematopoiesis are complex. Proposed mechanisms include reactivation of tumor suppressor genes, induction of Type I interferon signaling (*5, 6*), cytotoxicity against malignant clones (*7*), and inhibition of protein synthesis (by AZA, but not dAZA) (*8*). HMA response does not require clearance of mutated HSCs, and the increased circulating cells at response are in many cases the differentiated progeny of these mutated cells (*9–12*). Importantly, AZA may exert effects that are extrinsic to hematopoietic stem and progenitor cells (HSPC) by altering the function of non-malignant immune or stromal cells in the bone marrow (BM) (*13*).

The BM microenvironment contains multiple distinct cell types including leukocytes, erythrocytes, adipocytes, endothelial cells, fibroblasts, osteoblasts, and CXCL12-abundant reticular (CAR) cells that collectively support hematopoiesis (*14, 15*). Furthermore, dysregulation of the BM microenvironment has been increasingly implicated as a driver of malignant hematopoiesis (*16–20*); constitutive activation of β-catenin in osteoblasts causes AML in mice (*21*), while regulatory T cells have been linked to MDS outcomes (*22–24*). In MDS, the ratio of mature to immature NK cells is associated with clinical outcomes (*25*), and NK cell cytotoxic function is impaired (*26*). Furthermore, CXCL12 expression is elevated in bone marrow CAR cells in MDS (*27*). However, systematic changes in the BM microenvironment during HMA treatment have been poorly characterized to date.

Here we present a longitudinal, multi-modal analysis of higher risk (HR)-MDS patients receiving AZA monotherapy in a clinical trial setting (NCT03493646) (*11*). Clinical response was accompanied by an expansion of granzyme B (GzmB)^+^CD56^+^CD8 T cells, substantial changes in peripheral blood soluble factors, and spatial reorganization of the bone marrow microenvironment including increased CAR-cell neighborhoods. Together, these findings implicate both immune activation and stromal remodeling as correlates of AZA response in MDS.

## RESULTS

### CD8 T cell subsets expand when patients respond to AZA

HR-MDS patients who were enrolled in a multi-center phase 2 clinical trial (oral AZA; (NCT03493646)) received six cycles of injection AZA followed by six cycles of oral AZA (CC-486) (*11*). In each 28-day cycle (C), patients received seven daily injections of AZA (C1-C6) or 21 days of CC-486 (C7-C12) with response assessments at the end of six and 12 cycles of treatment. Clinical samples collected longitudinally at various time points were used for downstream analyses using multiple parallel platforms (Fig. 1A).

**Figure 1.**
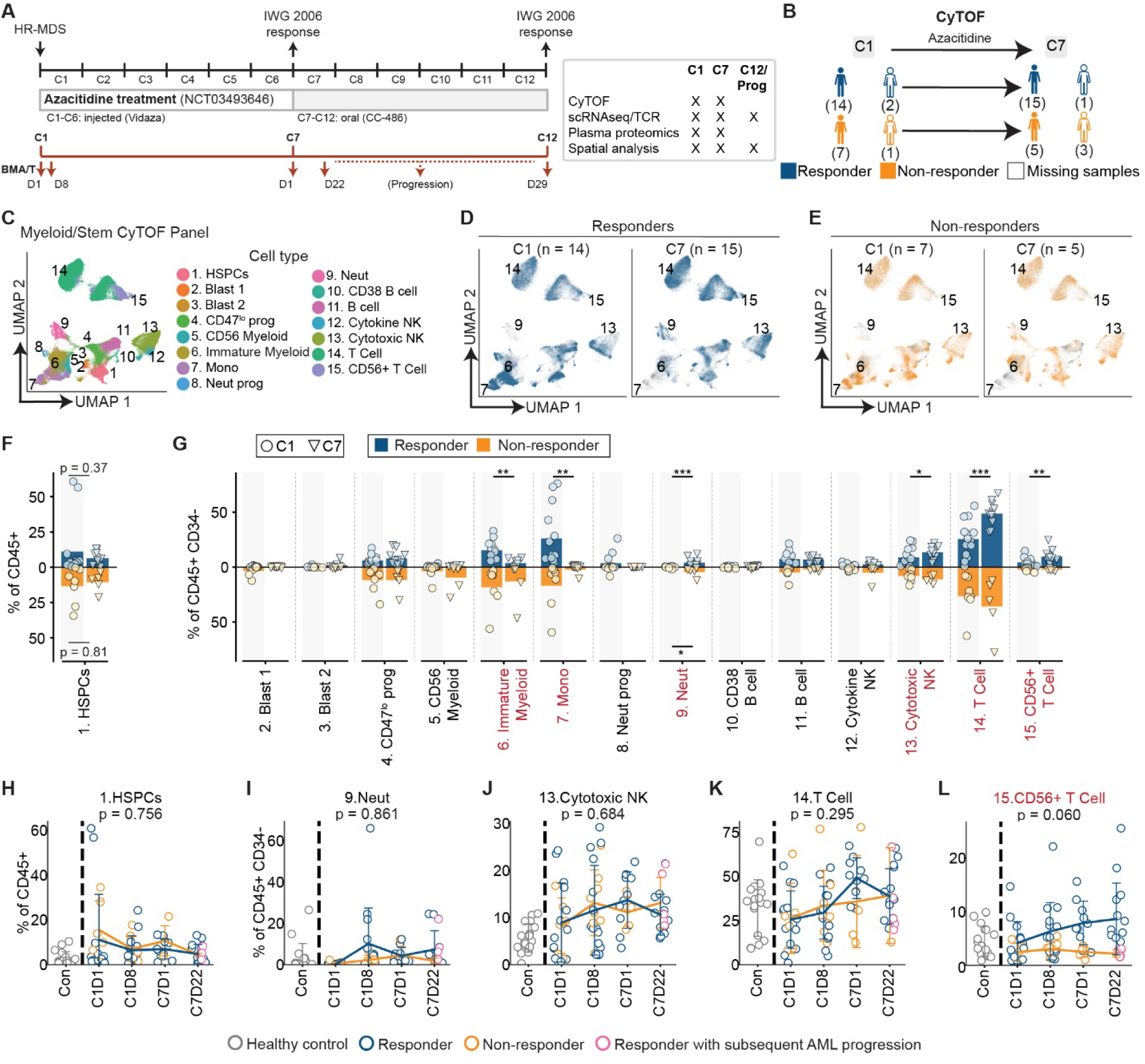
T cells expanded when patients responded to AZA. (**A**) Schematic showing AZA treatment schedule and sample collection timepoints (cycle (C), day (D), bone marrow aspirate + or – trephine (BMA/T)), and assays performed. (**B**) Schematic showing CyTOF patient cohort and clinical outcomes. (**C-E**) UMAP embeddings showing (**C**) cell types identified in human bone marrow, and changes between diagnosis and C7 in (**D**) responders, and (**E**) non-responders. (**F-G**) Changes in (**F**) CD34^+^ and (**G**) CD34^-^leukocyte frequency between diagnosis (C1) and C7. P-values: linear mixed-effects model, bar height shows the mean. (**H-L**) Longitudinal changes in abundance of (**H**) CD34^+^ HSPCs, (**I**) Neutrophils, (**J**) Cytotoxic NK, (**K**) total T cells, and (**I**) CD56^+^ T cells, grouped by clinical response. Pink markers indicate responders with subsequent disease progression. P-values: generalized linear mixed model, error bars show standard deviation. * p < 0.05, ** p < 0.01, *** p < 0.001.

We first used mass cytometry by time-of-flight (CyTOF) to characterize longitudinal changes in abundance of stem, lymphoid, and myeloid cells in BM aspirates (BMA) from a total of 24 patients at pre-treatment C1 day (D) 1 (n=21), C1D8 (n= 23), C7D1 (n=16) and C7D22 (n=17), with variation in numbers reflecting incomplete longitudinal sample availability (Fig 1B, Supplemental Table 1). A panel of 35 antibodies was used to profile BM cell populations (Supplemental Table 2). FlowSOM clustering identified the major leukocyte populations in human BM (Fig. 1C, Fig S1A), including CD34^+^ HSPCs, three myeloid progenitor cell clusters, monocytes, neutrophils, and lymphocyte populations comprising B cells, NK cells, and T cells. Two blast cell clusters were also identified, both expressing CD38, CD123, and moderate levels of CD33, with one cluster additionally expressing HLA-DR. Plasmacytoid dendritic cells and innate lymphoid cells were not detected likely due to their low abundance in human BM samples (*28, 29*).

We first compared changes in leukocyte frequencies between C1 and C7 in patients who achieved a clinical response to AZA after six treatment cycles (responders, Fig. 1D) and those who did not (non-responders, Fig. 1E). Overall, the frequency of HSPCs remained stable in both groups (Fig. 1F). To determine whether treatment altered specific HSPC subsets, we reclustered the CD34^+^ compartment. Seven clusters were identified (Fig. S1B, C), including two CD38low HSC clusters, one of which also expressed CD33, a marker of leukemic stem cells. Conventional progenitor populations, including megakaryocyte erythroid progenitors (MEPs), common myeloid progenitors (CMPs) and granulocyte monocyte progenitors (GMPs), were also identified, together with two atypical progenitor subsets expressing CD33 and HLA-DR but lacking CD123, one of which also expressed the adhesion molecule CD56 (Fig. S1C). Among these subsets, only GMPs increased significantly from C1 to C7 in responders, whereas no significant changes were observed in non-responders (Fig. S1D, E). No other HSPC subset showed significant temporal changes in frequency.

Both response groups showed reduced frequencies of monocyte and immature myeloid cells, although these changes reached significance only in responders, together with increased neutrophil frequencies (Fig. 1 G). In contrast, the frequencies of cytotoxic NK cells, T cells, and CD56^+^ T cells increased only in responders. We next compared the temporal trajectories of these cell populations across all sampling timepoints. Trajectories of HSPCs, neutrophils, cytotoxic NK cells, and total T cells did not differ between responders and non-responders (Fig. 1H, I, J, K). However, the CD56^+^ T cell cluster showed a strong trend towards increased abundance in responders (Fig. 1L) prompting a more detailed examination of T cell subset composition.

To further characterize changes in T cell populations, we used a second CyTOF panel comprising 33 antibodies enriched for T and NK cell markers (Supplemental Table 3). FlowSOM clustering identified multiple lymphocyte populations, including cytokine-producing and cytotoxic NK cells, regulatory T (Treg) cells, cytotoxic CD4 T cells, and four CD8 T cell subsets, (Fig. 2A, Fig. S2A). Discrete CD8 T cell clusters were distinguished by differential expression of the cytotoxic effectors GzmB and perforin, and CD56 (Fig. S2A). Comparison of cluster frequencies between C1 (Fig. 2B) and C7 (Fig. 2C) were consistent with findings from the initial CyTOF panel. HSPC frequencies remained unchanged (Fig. 2D), whereas both responders and non-responders showed decreased monocyte and increased neutrophil frequencies (Fig. 2E). In contrast to previous reports (*22–24*), Treg frequencies were unchanged in both groups. However, the frequency of GzmB^+^CD56^+^CD8 T cells increased in responders but not in non-responders (Fig. 2E). Longitudinal analysis across all sampling timepoints further showed that both GzmB^+^CD56^+^ and GzmB^+^CD56^-^CD8 T cell populations increased selectively in responders (Fig. 2F, G). Notably, patients who later progressed from MDS to acute myeloid leukemia after C7D22 had low frequencies of GzmB^+^CD56^+^CD8 T cells at C7D22 (Fig. 2F, pink circles). In contrast, the trajectories of cytotoxic CD4 T cells, cytotoxic NK cells, and Tregs did not differ between responders and non-responders (Fig. 2H, I, J).

**Figure 2.**
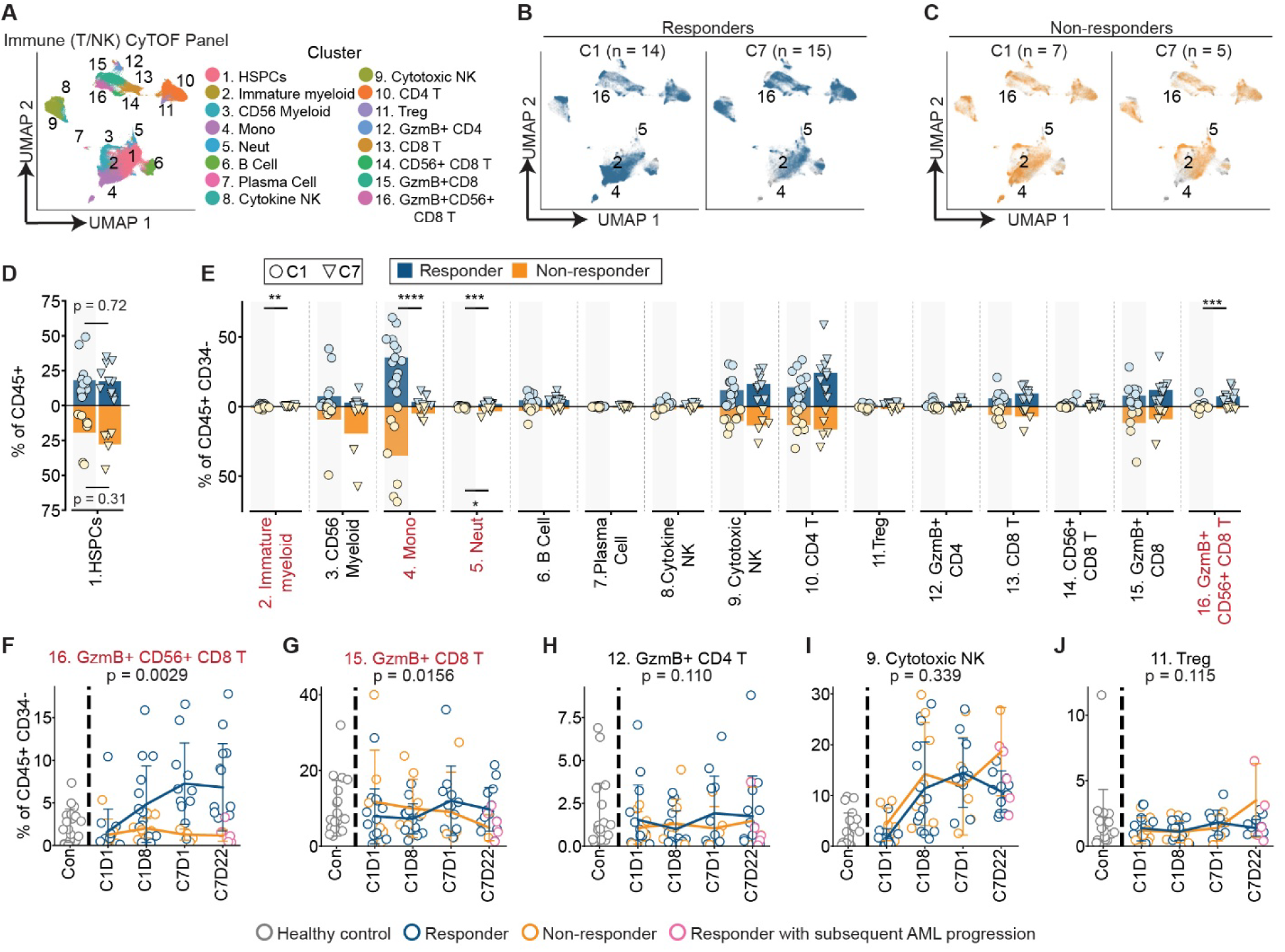
CD8 T cells expanded when patients responded to AZA. (**A-C**) UMAP embeddings showing (**A**) cell types identified in human bone marrow using a T/NK-focused antibody panel, and changes between diagnosis and C7 in (**B**) responders, and (**C**) non-responders. (**D-E**) Changes in (**D**) CD34^+^ and (**E**) CD34^-^ leukocyte frequency between diagnosis (C1) and C7. P-values: linear mixed-effects model, bar height shows the mean. (**F-J**) Longitudinal changes in abundance of (**F**) GzmB^+^CD56^+^CD8 T cells, (**G**) GzmB^+^CD8 T cells, (**H**) GzmB^+^CD4 T cells, (**I**) Cytotoxic NK cells, and (**J**) Tregs, grouped by clinical response. Pink markers indicate responders with subsequent disease progression. P-values: generalized linear mixed model, error bars show standard deviation. * p < 0.05, ** p < 0.01, *** p < 0.001.

To further resolve treatment-associated changes in CD8 T cell populations, we re-clustered the CD8^+^ compartment, identifying nine subsets including naïve/memory, memory, and an activated population expressing CD38, PD-1, and low-levels of GzmB. Four tissue-resident memory T cell populations were distinguished by differential expression of CD56 and CD127, whereas two cytotoxic GzmB^+^ perforin^+^ subsets were distinguished by CD56 expression (Fig. S2B, C). The CD56⁺GzmB⁺ subset expressed higher levels of CD57, perforin, GzmB, KLRG1, and CD16 than the CD56⁻ GzmB⁺ subset, consistent with an NK-like phenotype (Fig S2D). Comparison of C1 and C7 samples revealed modest increases in the naïve/memory, memory, and CD56^+^CD127^+^ Trm populations in responders, together with more pronounced expansion of both GzmB^+^ cytotoxic CD8 T cell subsets (Fig. S2E). Longitudinal analysis across all sampling timepoints similarly demonstrated differential trajectories of the naïve/memory, GzmB^+^CD56^-^, and GzmB^+^CD56^+^CD8 T cell populations between responders and non-responders (Fig. S2F). In contrast, the activated and Trm subsets showed no significant longitudinal differences between response groups.

Finally, we asked whether expansion of CD8 T cells was accompanied by changes in markers of activation or cytotoxicity. Longitudinal trajectories of CD38, CD69, HLA-DR, and PD-1 did not differ between responders and non-responders in either GzmB^+^ CD8 T cell subset (Fig. S2G, H), although GzmB^+^CD56^+^ cells in responders showed a modest trend towards higher expression of GzmB, CD57, and perforin, particularly at later timepoints (Fig. S2H). Collectively, these findings indicate that expansion of GzmB^+^CD8^+^ cytotoxic CD8 T cells is associated with clinical response to AZA, and occurs largely independently of changes in activation marker or effector molecule expression.

### IFN signaling is increased in responder T cells

We next performed single-cell RNA sequencing coupled with T cell receptor (TCR) sequencing on CD3^+^ BM T cells from a subset of patients (nine patients at three timepoints; C1, C7, and either C12 or progression based on sample availability; Fig. 3A). Clustering based on gene expression identified diverse CD4 and CD8 T cell populations (Fig. 3B). Within the CD4 compartment, we annotated naïve (*CCR7*, *LEF1*, *SELL*), cytotoxic (*GZMB*), and regulatory T (Tregs; *IL2RA*, *FOXP3*) cell populations. Within the CD8 compartment, three *GZMK*-expressing effector subsets were resolved based on differential expression of *MTRNR2L8*, *IL7R*, *GZMH*, *ITGB1*, and *CMC1*. Additional populations included a Trm cluster (*CD69*+, *KLF2*^lo^), and two *GZMB*-expressing terminal effector memory T cells (TEMRA) subsets. One TEMRA subset expressed *ZNF683* (HOBIT), whereas the other displayed an NK-like transcriptional program characterized by *KLRF1* and *FCGR3A* expression (Fig. 3B, C).

**Figure 3.**
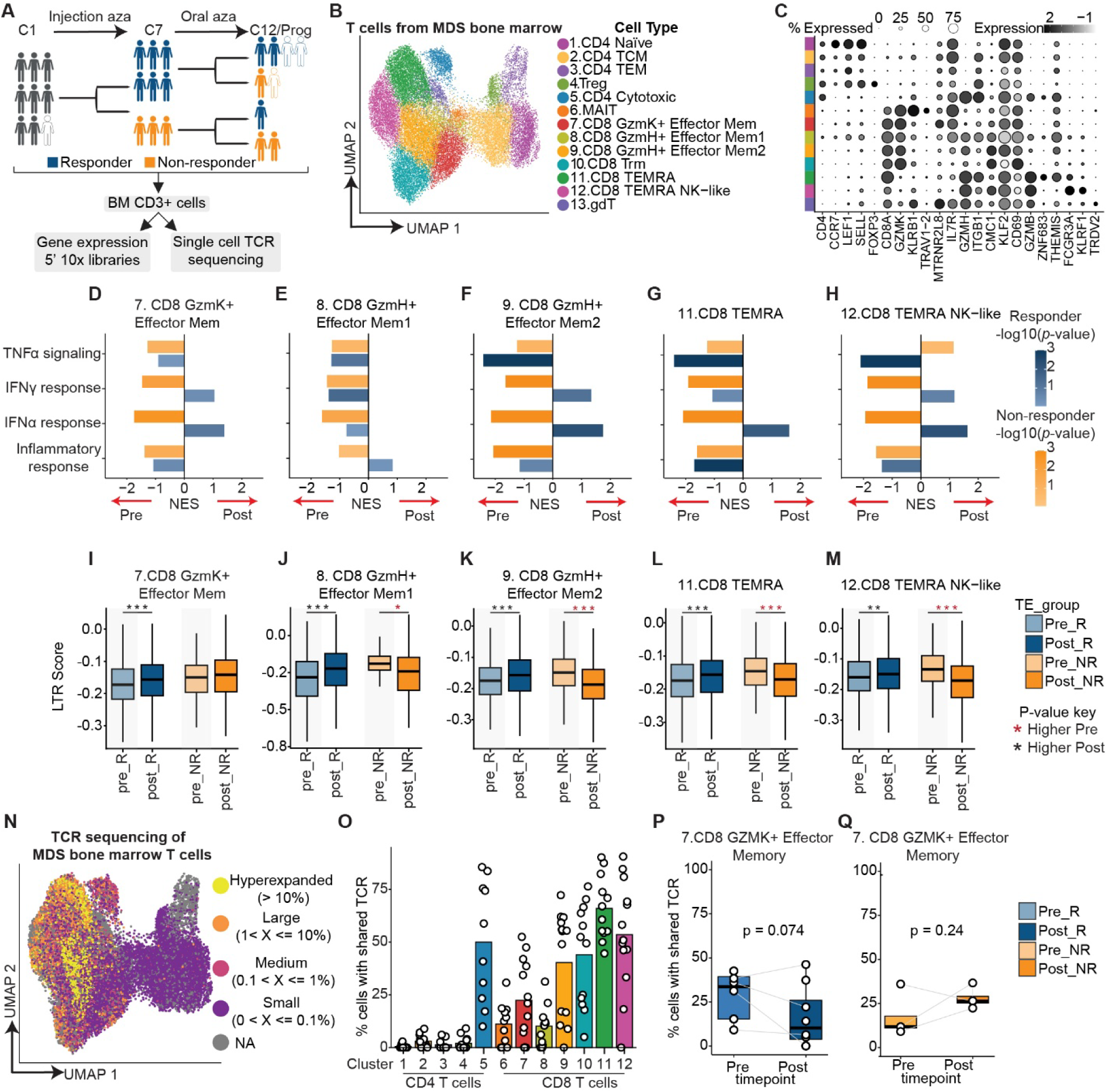
Increased IFNα signaling in BM T cells from patients who responded to AZA. (**A**) Schematic showing T cell single-cell sequencing cohort. Five healthy donors (not shown) were included in sequencing pools. (**B**) UMAP embedding of single-cell RNAseq bone marrow CD3^+^ T cells. (**C**) Expression of selected marker genes in the indicated T cell subsets. (**D-H**) Changes in pathway activity in selected T cell subsets pre– and post-AZA treatment in responders and non-responders. (**I-M**) Expression of LTR elements in selected T cell subsets pre– and post-AZA treatment. Asterisk color indicates the direction of change from pre– to post-treatment. (**N**) UMAP embedding colored by TCR clone size. (**O**) Percentage of expanded clones for each T cell subset. (**P-Q**) Comparison of clonality between pre– and post-AZA treatment in GzmK^+^ effector memory T cells in (**P**) responders, and (**Q**) non-responders. Paired t-test. * p < 0.05, ** p < 0.01, *** p < 0.001.

We next compared Hallmark pathway enrichment (*30*) in cytotoxic CD8 T cell subsets before and after AZA treatment. Interferon (IFN)-α signaling was generally increased in responders but decreased in non-responders across CD8 T cell populations (Fig. 3D, E, F, G, H). A similar pattern was observed for IFNγ signaling, although signaling decreased in responders within CD8 GZMH^+^ Effector Mem 1 and CD8 TEMRA subsets. In contrast, TNFα signaling and inflammatory response pathway activity generally decreased in both response groups, with the exception of the NK-like TEMRA population, in which TNFα signaling increased in non-responders. Together these findings indicate that clinical response to AZA was associated with selective remodeling of inflammatory signaling pathways in cytotoxic CD8 T cells, rather than a generalized inflammatory response.

AZA can induce transcription of transposable elements (TEs) and activate type I IFN signaling through viral mimicry (*5, 6*). We therefore asked whether altered TE or IFN gene expression could explain the elevated IFNα (type I IFN) gene signature observed in responder CD8 T cells. Long terminal repeat (LTR) element expression increased after AZA treatment in cytotoxic CD8 T cells from responders but decreased in non-responders in three of four clusters (Fig. 3I, J, K, L, M). In contrast, short interspersed nuclear element (SINE) and long interspersed nuclear element (LINE) expression showed variable, cluster-specific changes (Fig. S3A, B, D, E). To determine whether cytotoxic CD8 T cells were a direct source of type I IFNs, we examined interferon transcript expression. Type I IFN transcripts were undetectable in T cells. Although IFNγ (type II) and type III IFNs can also induce interferon-stimulated genes (ISGs), *IFNG* expression showed only a modest decline after AZA treatment in responder effector memory T cell subsets (Fig. S3F), whereas *IFNL1* expression was unchanged (Fig. S3G). As single cell RNA sequencing data from CD34^+^ cells from the same patient cohort were available (*31*), we next examined HSPCs as a potential source of IFNs. As in T cells, IFN transcripts were largely undetectable, although *TGFB1* was readily detected, confirming data quality (Fig. S3H). TE expression showed modest decreases in SINE, LINE, and LTR elements in responders but increased LINE and LTR expression in non-responders (Fig. S3I, J, K). Consistent with these findings, bulk RNA sequencing of CD34^+^ cells from HR-MDS patients (*11*) revealed negligible expression of IFN transcripts (Fig. S3L). Collectively, these data suggest that the elevated type I IFN signaling observed in cytotoxic CD8 T cells was unlikely to result from autocrine IFN production by T cells or CD34^+^ progenitors and instead reflected IFN derived from other cell types within the bone marrow microenvironment.

We then investigated changes in the TCR repertoire following AZA treatment. T cell clones were classified as small (≥ 0.01– < 0. 1%), medium (≥ 0. 1 – < 1%), large (≥ 1 – < 10%), or hyperexpanded (≥ 10%), based on the number of cells sharing the same TCR sequence. Hyperexpanded clones were observed exclusively within the CD8 compartment, whereas large and medium clones were found in cytotoxic CD4 T cells and multiple CD8 T cell clusters (Fig. 3N, O). In contrast, the remaining CD4 T cell subsets consisted predominantly of cells with unique TCR sequences. These findings are consistent with previous reports that clonally expanded T cells in the BM of AML patients (*32*) and in peripheral blood of older individuals (*33*) are enriched within the CD8 compartment. We next examined changes in clonal expansion over time. In responders, the proportion of cells belonging to expanded clones tended to decrease within the *GZMK*-expressing effector memory population (Fig. 3P), whereas a trend toward increased clonal expansion was observed in non-responders (Fig. 3Q). No significant longitudinal changes in clonal expansion were detected in the remaining CD8 T cell populations.

Finally, we queried VDJdb and the Immune Epitope Database (IEDB) to identify TCRs predicted to recognize known neo-antigens or tumor-associated antigens (*34, 35*). Most annotated clonotypes were specific for viral antigens, including cytomegalovirus (CMV), Epstein-Barr virus (EBV), and influenza A (Fig S4A), whereas only a small number recognized self– or tumor-associated antigens including WT1, MLANA, BST2, IGF2BP2, and TYR (Fig. S4B, C). Across all patients, approximately 0.5% and 1% of TCRs mapped to self– or tumor-associated clonotypes in VDJdb and IEDB, respectively (Fig. S4D). We did not detect TCRs recognizing the CLK3 neo-junction reported to arise from SRSF2 splicing mutations (present in P01, P02, P08, P11 (*11*)) (*36*). Overall, these findings indicate that TCRs with known tumor antigen specificities are rare in this cohort and that the antigen specificities of the vast majority of TCR clonotypes remain undefined.

### Plasma proteomic profiles are profoundly altered in responders compared to non-responders

Cytokines and growth factors are important regulators of hematopoiesis and leukocyte function, and may contribute to the type I IFN gene signature observed in responder CD8 T cells (*37*). To define soluble factor changes associated with AZA treatment and clinical response, we used a multiplexed immunoassay (NULISA) (*38*) to quantify relative concentrations of 247 analytes in BM and peripheral blood (PB) plasma collected before and after treatment from 17 patients (Fig. 4A). Approximately 80% of analytes were detected in BM plasma and more than 90% in PB plasma (Fig. S5A). Several soluble factors differed between responders and non-responders before treatment. In BM plasma, CCL28, GFAP, and IL-36A were more abundant in responders, whereas TIMP1, CRP, CD80, and CXCL12 were enriched in non-responders (Fig. 4B). Similarly, PB plasma from responders showed higher concentrations of IRAK4 and CSF3, whereas CCL3, IL2RA, CHI3L1, IL-6, and CXCL12 were elevated in non-responders (Fig. 4C).

**Figure 4.**
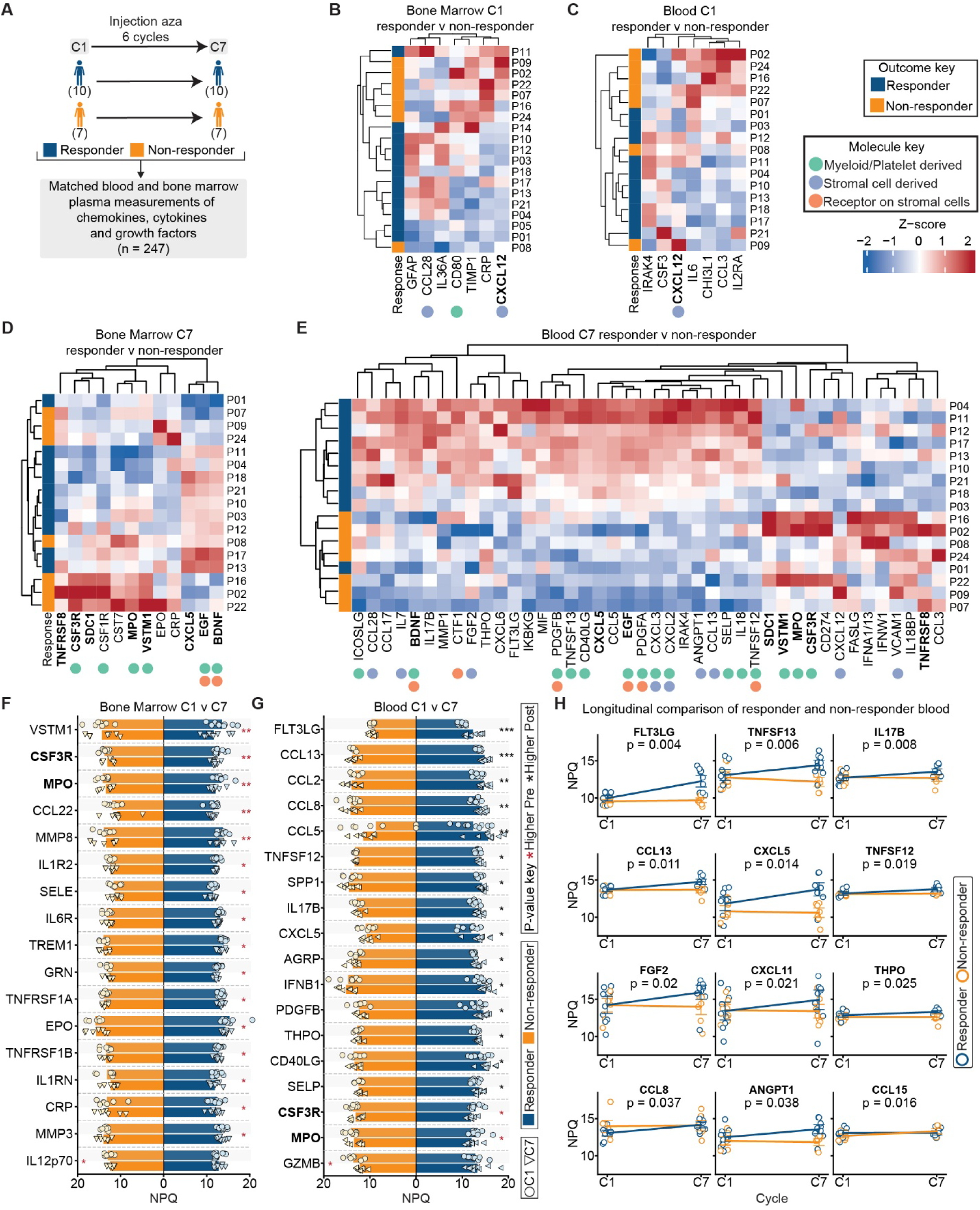
Significant alterations in abundance of soluble mediators when patients responded to AZA. (**A**) Schematic showing plasma proteomics cohort. (**B-E**) Heatmaps showing the relative abundance of soluble mediators at diagnosis (C1) in (**B**) bone marrow, and (**C**) blood plasma, and at C7 in (**D**) bone marrow, and (**E**) blood plasma. (**F-G**) Changes in soluble mediators in (**F**) bone marrow, and (**G**) blood plasma at diagnosis and C7. Bar height indicates the mean, asterisk color indicates the direction of change from pre– to post-treatment. P-values: linear mixed-effects model. (**H**) Longitudinal change in selected soluble mediators, grouped by clinical response. P-values: generalized linear mixed model, error bars show standard deviation. *p < 0.05, **p < 0.01, ***p < 0.001.

Differences between responders and non-responders were more pronounced at the C7 clinical response assessment, with 12 and 42 differentially abundant analytes detected in BM and PB plasma respectively, including eight shared between the two compartments (Fig. 4D, E, Supplemental Table 4). BDNF, EGF, and CXCL5 were increased in both BM and PB plasma from responders, consistent with improved platelet production. Several additional factors elevated in responder PB, including PDGFA, PDGFB, and FGF2, are associated with hematopoietic recovery but also signal through receptors expressed by bone marrow stromal populations, including CXCL12-abundant reticular (CAR) cells, fibroblasts, and vascular smooth muscle cells (*39*), raising the possibility that they contribute to hematopoietic recovery through effects on the BM niche. In contrast, plasma CSF1R, CSF3R, TNFRSF8, SDC1, VSTM1 and MPO were elevated in both BM and PB from non-responders. As these proteins are predominantly cell surface or intracellular molecules, their elevated concentrations likely reflect ongoing hematopoietic cell death, rather than active secretion. This interpretation is supported by persistently elevated CRP (a protein produced by the liver that binds microbes or damaged/dying cells (*40*)) in BM plasma from non-responders at both C1 and C7, but not PB. Soluble CSF1R and CSF3R may also contribute to dysplasia by acting as ligand traps that impair monocyte and neutrophil maturation.

Unexpectedly, CXCL12 concentrations were higher in non-responders than responders in BM and PB plasma at C1 and remained elevated in PB at C7. Given its established role in retaining CXCR4^+^ haematopoietic cells within the bone marrow (*41*), elevated CXCL12 may represent both a biomarker of AZA nonresponse and a potential contributor to MDS pathogenesis by promoting stem/progenitor cell retention or impairing normal hematopoietic differentiation and trafficking.

We next examined plasma ratios between selected cytokines and their inhibitors or soluble receptors, reasoning that soluble receptors can function as ligand traps (e.g. (*42*)). The ratios of IL-18 to its negative regulators, IL-18BP and soluble IL18R1, were higher in PB plasma from responders at C7 than in non-responders (Fig. 4E, Fig. S5B), consistent with increased IL-18 bioavailability. IL-7 concentrations were also elevated in responder PB at C7 (Fig. 4E). Together, these findings are consistent with enhanced cytokine signaling that may support expansion of GzmB^+^CD56^+^CD8^+^ T cells in responders. Similarly, the ratio of plasma CD40LG (predominantly from platelets and T cells (*43*)) to CD40 and the ratios of myeloid-derived TNFSF13 (APRIL) to its receptors TNFRSF17 (TACI) or TNFRSF13B (BCMA) were increased in responders (Fig. 4E, Fig. S5B). APRIL and APRIL ratios also increased from C1 to C7 in responders (Fig. S5C) but not in non-responders (Fig. S5D). As CD40LG and APRIL promote B cell and plasma cell survival (*44, 45*), these findings are consistent with enhanced humoral immune support in responders. Note that plasma cells were not represented in our CyTOF analysis because of their low CD45 expression.

We next compared concentrations of soluble factors at C1 vs C7 within responders or non-responders (Fig 4F, G). In BM plasma (Fig. 4F), 16 analytes, including CSF3R and MPO and the inflammatory markers IL1-R2, IL1-RN, TNFRSF1A and TNFRSF1B decreased significantly from C1 to C7 in responders, consistent with a reduction of dysplastic progenitors. In contrast, only IL-12p70 (mainly produced by antigen presenting cells to induce Th1 type immunity and support cytotoxic T cells) (*46*)) changed significantly in non-responders, decreasing between C1 and C7, consistent with limited support for cytotoxic T cell responses. A similar pattern was observed in PB plasma (Fig. 4G). Responders again showed reduced MPO and CSF3R together with increased concentrations of 15 soluble factors, including FLT3LG, a regulator of myeloid progenitor proliferation and differentiation (*47*). In contrast, GzmB was the only analyte that changed significantly in non-responders, decreasing after treatment.

Comparing longitudinal trajectories of responders with non-responders there were no differences in BM (not shown), but many molecules showed greater increases in responders compared with non-responders in blood (Fig. 4H) including the chemokines CCL13, CXCL5, CXCL11 and CCL8; factors regulating hematopoiesis: FLT3LG, ANGPT1 and THPO; and cytokines TNFSF12 (TWEAK) and TNFSF13 which were all increased in responders. Only one molecule, CCL15, showed a longitudinal increase in non-responders compared with responders, but the change was modest.

To examine how soluble factor abundance relates to disease burden and immune activity, we computed Spearman correlations between plasma protein abundances and frequencies of blasts or CD8 T cell subsets. Increasing blast burden was associated with reduced IL-12B and IFNL2/3 and increased levels of the soluble receptors CSF1R, SIRPA, IL-6R, IL-7R, IL-1RL1, and IL-1R2 (Fig. S4E). In contrast, expansion of GzmB^+^CD56^+^CD8 T cells was positively associated with IL-12p70 and IL-12B thus identifying IL-12 is a candidate regulator of cytotoxic CD8 T cell expansion in MDS patients. CSF3 abundance negatively correlated with cytotoxic CD8 T cell frequency, whereas IL-15 was negatively associated with GzmB⁺CD56⁻ but not GzmB⁺CD56⁺ CD8 T cells, suggesting that expansion of the latter population occurs independently of IL-15 (Fig. S4F). Together, these findings indicate that clinical response to AZA was accompanied by coordinated changes in soluble plasma factors that are associated with reduced disease burden, reinvigoration of the bone marrow stromal niche and improved hematopoiesis, and expansion of cytotoxic CD8 T cells.

### AZA treatment induces architectural changes in the BM microenvironment

AZA-response occurs in the context of 3-dimensional tissue structure; information lost from aspirate specimens. Furthermore, some cell types (e.g. CAR stromal cells) are poorly represented in aspirate samples (*48*). We therefore used multiplexed immunofluorescent imaging of fixed BM trephine sections to explore spatial relationships of HSPCs and T, CAR and endothelial cells (Fig. 5A). Nuclei were marked by DAPI. T cell populations were identified using CD3 plus CD4 or CD8, while CXCL12 was used to mark CAR cells and CD34 was used to identify HSPCs and endothelial cells; endothelial cells were distinguished from HSPC by morphology (Fig. 5B). Nucleated cells not assigned to a defined population (“unknown” cells) were also quantified to capture differences in BM cellularity. Ki67 and PD1 were included as markers of proliferation and T cell exhaustion, respectively.

**Figure 5.**
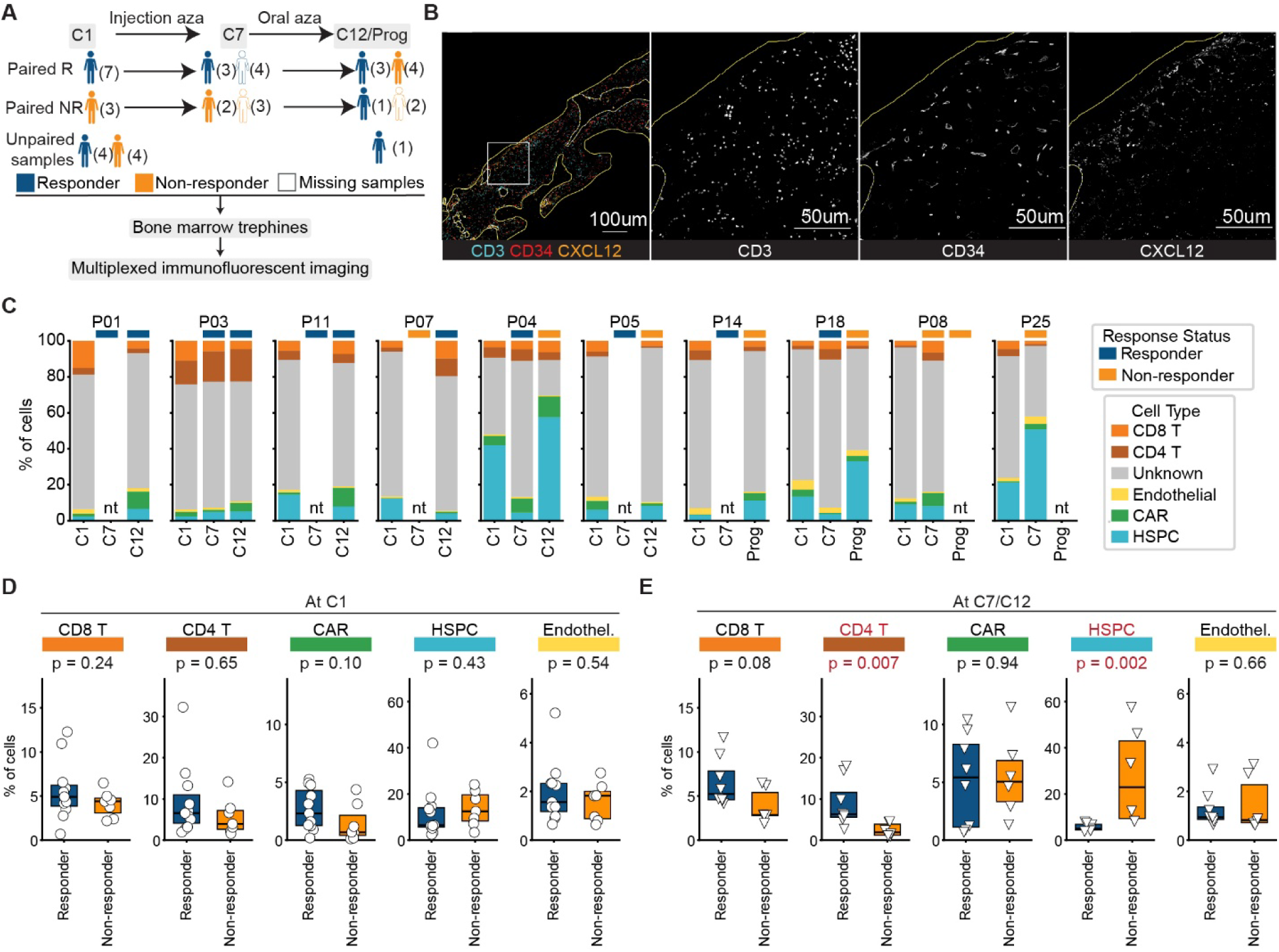
Elevated CD4 T cells in bone marrow when patients responded to AZA. (**A**) Schematic showing trephine cohort. (**B**) Representative multiplexed immunofluorescence image of a bone marrow trephine section. (**C**) Longitudinal cell type distribution in patients with paired pre– and post-AZA trephine specimens available. nt: no trephine at this timepoint. (**D-E**) Percent of total cells belonging to each tracked cell type at (**D**) diagnosis (C1), and (**E**) C7 or C12. P-values: Mann-Whitney U-test. *p < 0.05, **p < 0.01, ***p < 0.001.

We first examined changes in cell frequencies in the available paired samples. Four responders (P01, P03, P11, and P07) showed only minor changes in the frequencies of cells in immunophenotyped trephines comparing diagnosis (C1) to the timepoint(s) of response (C7 or C12; Fig. 5C). Two patients who initially responded and then progressed (P04 and P18) showed a substantial decrease in trephine HSPCs at clinical response (C7), followed by a rebound at progression (C12; Fig. 5C). One patient who never responded (P25) also showed a large increase in HSPC frequency from diagnosis (C1) to non-response C7 (Fig. 5C). In three patients where only diagnosis (C1) and a non-response timepoint were available (P05, P08, and P14), there were only minor changes in cell frequency between C1 and C7 or C12 (Fig. 5C). Overall, these findings confirm that the imaging panel reliably detects largely expected changes in HSPC frequency.

To determine whether baseline cell type abundance could predict clinical response, we compared cell type frequencies at diagnosis between responders and non-responders. No significant differences were observed at diagnosis, although CAR cells tended to higher frequency in responders (Fig. 5D). At the clinical assessment timepoint, HSPCs were more frequent in non-responders, while CD4 T cells were more frequent in responders, and CD8 T cells also showed a trend towards higher frequency in responders (Fig. 5E). Frequency of CAR cells, Ki67 expression in HSPCs and T cells, and PD-1 expression on T cells did not differ between groups at either timepoint (Fig. 5D, E, Fig. S6).

We also assessed differences in local cellular neighborhoods. To define neighborhoods, each annotated CD8 T, CD4 T, endothelial, CAR or HSPC cell was individually treated as a focal cell, for which distance to all neighboring cells within a 100 μm radius was measured and weighted via an exponential decay function to produce a proximity weighted neighborhood matrix for each focal cell (Fig. 6A). Leiden clustering was then used to identify 59 neighborhood clusters, (“neighborhoods”), which were visualized on a UMAP plot (Fig. 6B). Neighborhoods 45 and above, which contained fewer than 1,000 cells, were excluded from further analysis (Fig. S7A, Supplemental Table 5).

**Figure 6.**
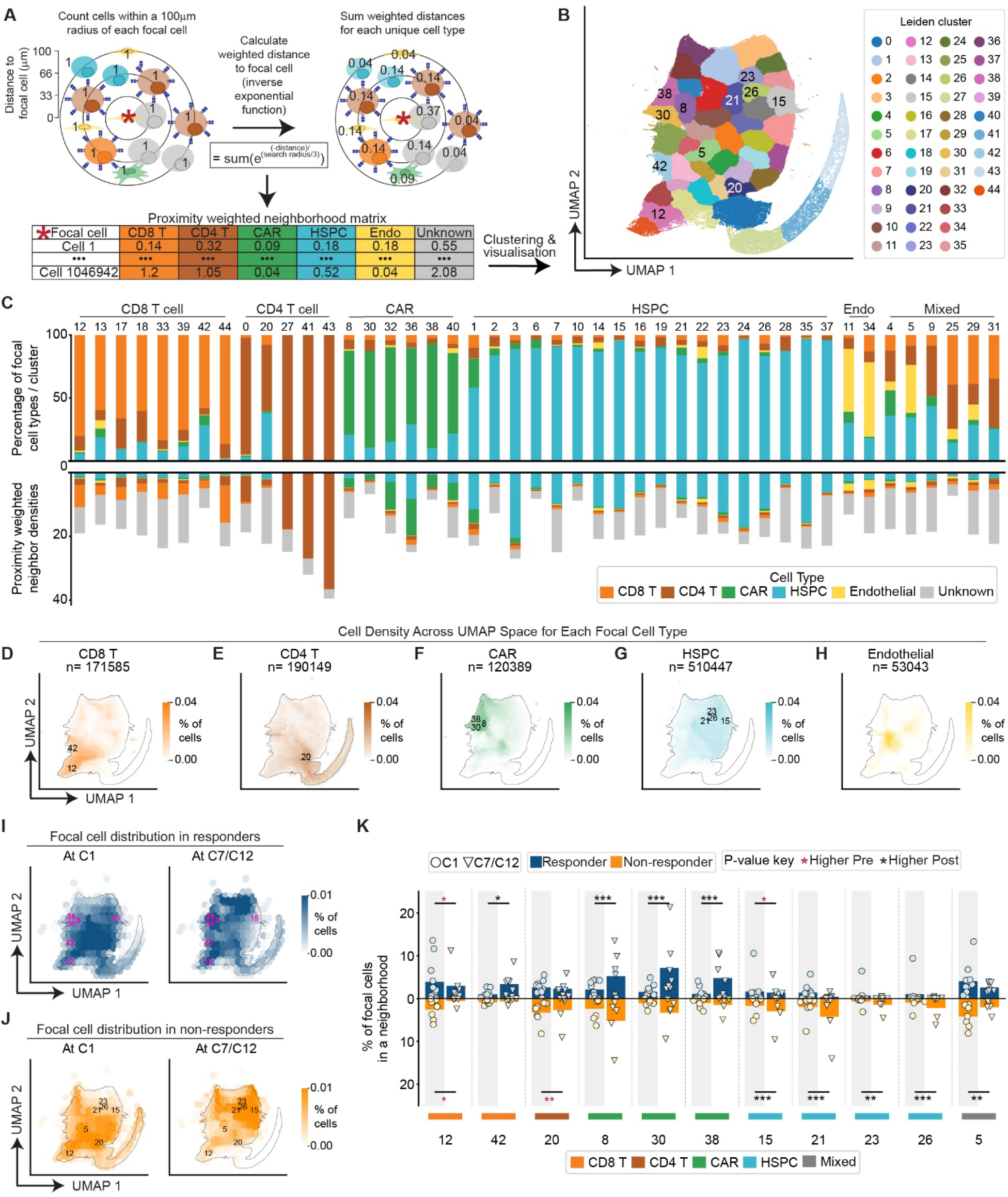
Specific CAR cell-rich/HSPC-depleted and CD8 T cell-rich neighborhoods expanded in responder BM. (**A**) Schematic showing method to define a proximity-weighted neighborhood matrix of cells within 100µM of each annotated focal cell. (**B**) UMAP embedding highlighting clustered neighborhoods. (**C**) Percentage of focal cells contributed by each annotated cell type in each clustered neighborhood (top) and proximity-weighted densities of non-focal neighboring cells (bottom) plotted by calculating then stacking the averaged sum of each column for all rows representing a neighborhood cluster in the proximity-weighted neighborhood matrix (see **(A)**). (**D-H**) UMAP embeddings highlighting distributions of focal cells; (**D**) CD8 T cells, (**E**) CD4 T cells, (**F**) CAR cells, (**G**) HSPCs, (**H**) endothelial cells. (**I-J**) UMAP embeddings showing distributions of focal cells at diagnosis and clinical response in (**I**) responders, and (**J**) non-responders. Dotted line in (**D-J**) indicates the approximate boundary of the UMAP embedding. (**K**) Frequencies of focal cells in each neighborhood at diagnosis and at response or non-response. Bar height indicates the mean, asterisk color indicates the direction of change from C1 to C7. P-values: linear model with treatment response as a fixed effect; *p < 0.05, **p < 0.01, ***p < 0.001.

All neighborhoods contained multiple annotated cell types, except CD4 T cell rich neighborhoods 27, 41, and 43 (Fig. 6C). Most neighborhoods were present across the majority of specimens, but a few were restricted to a small subset (Fig. S7A); these tended to have low Shannon diversity (Fig. S7A), indicating that the bulk of cells were contributed by a disproportionately small number of trephines. CD8 T cells (Fig. 6D), CD4 T cells (Fig. 6E), CAR cells (Fig. 6F), and HSPCs (Fig. 6G) were distributed throughout the focal cell UMAP, suggesting these cell types all reside within diverse local microenvironments, as expected. Nonetheless, regions of focal cell type enrichment were clearly evident (Fig 6D-H).

We next compared neighborhood frequencies between responders and non-responders. At diagnosis (C1), neighborhood frequencies and overall cell organization were similar between the two groups (Fig. 6I, J). However, at the time of clinical assessment (C7 or C12), responders had a higher proportion of cells within neighborhoods enriched for CD8 T cells (12, 13, 17, 18, 33, 42;

Fig. S7B, C), CD4 T cells (0, 20; Fig. S7B, C) or mixed cell populations (4, 9, 25, 29, 31; Fig. S7B, C). In contrast, non-responders had a higher proportion of cells within HSPC dominated neighborhoods (1, 7, 14, 15, 21, 22, 23, 26; Fig. S7B, C). Four neighborhoods enriched for CAR focal cells (8, 30, and 38) contained a higher proportion of responder cells, whereas one CAR-focal cell neighborhood (36) contained a higher proportion of non-responder cells (Fig. S7B, C). Clusters enriched for endothelial focal and proximal cells (11, 34; presumably blood vessels) did not differ between groups.

Finally, we compared neighborhood composition between diagnosis and clinical response assessment within each response group. In responders, the proportion of cells assigned to neighborhood 42, enriched for interactions between CD8 T cells and HSPC, and neighborhoods 8, 30, and 38, CAR cell neighborhoods depleted of HSPCs, increased (Fig. 6K). In contrast, a higher proportion of cells in non-responders were assigned to HSPC-rich neighborhoods (15, 21, 23, and 26), consistent with blast-enriched regions (Fig. 6K). Neighborhoods 12, enriched for CD8 T cells, and 20, uniquely enriched for CD4 T cells and HSPCs, contracted in non-responders. Taken together, these findings associate clinical response to AZA with reduced HSPC (or blast) self-association, increased spatial interactions between HSPCs (or blasts) and CD8 T cells, and remodeling of CAR cell organization within the bone marrow.

## DISCUSSION

Clinical response to HMAs is defined by improved circulating blood counts and decreased blasts and does not require clearance of mutated HSPCs or their progeny (*9–12*). HSPC intrinsic effects of AZA are reasonably well characterized, but even though HMAs might affect other cell types both directly and indirectly, comprehensive studies of HSPC-extrinsic responses to AZA are lacking. Here we tracked longitudinal changes in leukocyte frequency, cytokine abundance, and spatial organization of the BM microenvironment in patients treated with AZA monotherapy in a clinical trial setting and found selective expansion of GzmB^+^CD56^+^CD8^+^ T cells, altered inflammatory signaling in specific T cell subsets, a profoundly altered cytokine milieu, and remodeling of BM spatial organization in patients who respond to AZA therapy.

The role of CD8 T cells in MDS is unclear: cytotoxicity may be impaired or excessive (*49, 50*), azacitidine may increase TCR diversity (*51*), and CD8 T cells may be more differentiated in lower-risk (LR-MDS) compared to healthy controls (*52*). An elevated frequency of cytotoxic CD8 T cells at diagnosis has also been associated with unfavorable outcomes (*53*). Here we found an expansion of GzmB⁺CD56⁺CD8⁺ T cells in BM of AZA responders (Fig. 2), alongside an increase in the frequency of the BM neighborhood most enriched for associations of CD8 T cells with HSPC in responders (Fig. 6, neighborhood 42). CD56^+^CD8 T cells have been reported to produce cytokines in a TCR-independent manner (*54*), implying they can respond to innate signals such as IL-12 and IL-18 (a cytokine which was higher in PB of responders in our cohort) independent of cognate antigen. We observed only modest differences in T cell clonality between pre– and post-treatment samples, and other studies reported that BM T cell clonality did not change longitudinally within responders or non-responders (*55*) and we did not observe expansion of TCRs typical of innate-like CD56^+^ T cells known to recognize non-protein antigens (*56*). T cell clonality was reported as higher in MDS BM compared to healthy controls (*55*) and higher TCR diversity in the blood was associated with longer event-free survival in AML patients receiving AZA (*57*). Our data is consistent with cytotoxic CD8 T cells acting in responders via innate rather than TCR-mediated signaling.

We observed multiple changes in inflammatory signaling pathways in specific T cell subsets. Type I IFN signaling increased in responders and decreased in non-responders across cytotoxic CD8 T cell subpopulations. This finding is consistent with previous reports that IFNα signaling was elevated in many clusters of CD8 T cells in both HR-MDS and AML patients who respond to AZA treatment (*53*). Although LTR expression was increased in responder T cells and LTR elements are known to activate innate cytosolic dsRNA sensing and type I IFN signaling (*5, 6*), we did not observe increased IFN production in T cells or CD34^+^ cells implicating other cell types in the bone marrow microenvironment as the source. IFNγ signaling also increased in responders and decreased in non-responders across cytotoxic CD8 T cell subpopulations, consistent with a previous report of increased *IFNG* expression in CD8 TEMRAs of HR-MDS patients with long-term response to decitabine and PD-L1 blockade (*58*). TNFα signaling decreased in both responder and non-responder cytotoxic CD8 T cell subpopulations in our cohort, in contrast to a previous observation of elevated TNFα signaling in GzmK^+^CD8^+^ T cells in relapsed/refractory AML patients who responded to AZA + PD1 blockade (*59*), and an opposing report of elevated TNFα in CD8 T cells of non-responders (*53*). Although immune checkpoint blockade has not improved outcomes in unselected patients with HR-MDS (*60, 61*), our findings suggest that this warrants further investigation.

Our analysis of chemokines, cytokines, and growth factors revealed marked remodeling of the soluble proteome at the time of clinical response, particularly in peripheral blood. Several factors elevated in responders, including the platelet-derived growth factors PDGFA and PDGFB, likely reflect restoration of productive hematopoiesis rather than initiating the response. However, because these ligands signal to multiple bone marrow stromal populations, including CAR cells, they may also reinforce hematopoietic recovery through feed-forward interactions within the bone marrow niche. Most of the longitudinal changes represented increases in soluble factors in responders relative to non-responders. Among these, FLT3LG, produced primarily by stromal and activated T cells (*39, 62*)) and CD40LG (derived predominantly from activated platelets and CD4 T cells, (*43*)), were elevated in responder PB plasma at C7. Increased FLT3LG, which is required for granulocyte and dendritic cell development (*63*), may also partly explain improved PB counts. IL-7 and IL-18 (which can drive T cell proliferation) were higher, and IL-18BP (an inhibitor of IL-18) was lower, in PB of responders. Active IL-18 is cleaved and released by macrophages and dendritic cells in an inflammasome-dependent manner (*64*), while IL-18BP is produced by myeloid and endothelial cells (*65*). The elevated IL-18/IL-18BP ratio, alongside elevated IL-7, could contribute to the observed expansion of cytotoxic CD8 T cell subsets. Furthermore, IL-18 can drive the expression of perforin and GzmB in CD8 T cells (*66*) and is being incorporated into CAR T cell design to aid clearance of lymphoma (*67*) and AML (*68*). In contrast, CXCL12 was consistently elevated in non-responders in both bone marrow and peripheral blood plasma at diagnosis. Produced primarily by CAR cells, endothelial cells, and fibroblasts (*39*), CXCL12 regulates retention of CXCR4^+^ HSPCs and specific leukocyte subsets within the BM. Consistent with this role, pharmacological CXCR4 inhibition mobilizes HSPCs (*69*), elevated expression of CXCR4 on CD34^+^ cells is associated with reduced survival in MDS (*70*), and CXCL12 signaling can promote survival of MDS progenitors through induction of BCL2 (*71*). Together, these findings raise the possibility that excessive CXCL12 signaling contributes to disease persistence by promoting retention and survival of blasts while limiting productive hematopoiesis. These data also identify circulating CXCL12 as a candidate biomarker for predicting response to AZA and support further investigation of the CXCL12–CXCR4 axis as a therapeutic target in HR-MDS.

Simple counting of cell populations in BM trephine sections revealed few differences in relative cellular abundance other than the expected increases in HSPC blast frequencies in non-responders. In contrast, proximity-weighted neighborhood analysis identified marked differences in the spatial organization of the bone marrow microenvironment during treatment. HSPC-enriched neighborhoods remained largely stable in responders but expanded in non-responders. Conversely, responders exhibited increased representation of the neighborhood most enriched for HSPC/blast–CD8 T cell interactions (neighborhood 42), together with CAR-cell-enriched neighborhoods depleted of HSPC/blasts (neighborhoods 8, 30 and 38). These spatial changes are consistent with a model in which cytotoxic CD8 T cells contribute to hematopoietic recovery, either by eliminating dysplastic progenitors or by secreting cytokines that promote HSPC differentiation or function. Expansion of T cell neighborhoods could be supported by soluble factors elevated in responders, including CCL5, CCL13, IL-7, and ICOSLG, which regulate T cell recruitment, proliferation, and activation. Similarly, AZA-induced improvements in blood counts and production of growth factors that can signal to stromal cells (e.g. PDGFA, PDGFB, and BDNF, produced by platelets or megakaryocytes) may promote remodeling of the stromal compartment by acting on CAR cells and other bone marrow stromal populations. Together with our CyTOF, single-cell transcriptomic, and plasma proteomic analyses, these findings support a model in which azacitidine-induced hematopoietic recovery, cytotoxic CD8 T cell expansion, and remodeling of the CAR cell niche reinforce one another to establish a bone marrow microenvironment that favors productive hematopoiesis.

A key strength of this work is its basis in real world data collected with the consistency imposed by a formal and independently monitored clinical trial. Although validation of our findings in an independent clinical cohort would be ideal, longitudinal clinical sampling at fixed timepoints is rarely performed on patients receiving standard of care therapy, and comparable samples are difficult to source. Further mechanistic validation presents a particular challenge given the lack of suitable experimental models available. MDS patient-derived xenografts (*72*) do not replicate all aspects of human immunity and are not suitable to evaluate stromal cell response given their engineered BM microenvironment, while immunocompetent murine models are currently unable to recapitulate the molecular heterogeneity of the human disease. BM organoids (*73*) may provide a tractable model to explore the intrinsic effects of AZA on HSPCs in parallel with HSC-extrinsic effects mediated by immune or stromal cells. However, experimentation to validate how faithfully organoids engrafted with MDS HSPC, immune and stromal cells can recapitulate *in situ* MDS is required before such a model can be deployed widely.

## MATERIALS AND METHODS

### Study Design and Sample collection

Human BM and PB samples were collected as part of an open label phase 2 multicentre investigator-initiated trial (*11*) (NCT03493646: Evaluating In Vivo AZA Incorporation in Mononuclear Cells Following Vidaza or CC-486). All participants provided written informed consent for clinical research including genetic testing and collection/storage of human tissue. Use of samples received ethical approval (2022_ETH00727) from the South Eastern Sydney Local Health District Human Research Ethics Committee. Longitudinal samples were available for 25/40 patients). Precise sample size for each component of this study was dictated by specimen availability and practical experimental limitations and are detailed in Supplemental Table 1.

### Processing of BM aspirates and PB

BM and PB samples were collected and processed as described (*11*). Briefly, BM aspirates were diluted five-fold in RPMI 1640 (Thermo Fisher), and mononuclear cells (MNCs) isolated using Lymphoprep (ELITech Group). A subset of MNCs (previously depleted of CD34^+^ cells) were stained with anti-CD3 beads magnetic beads (Miltenyi) and CD3^+^ cells recovered using AutoMACS with program possel. MNCs were frozen in cryovials in 90% FBS + 10% DMSO and selected CD3^+^ cells were frozen in cryovials with fetal bovine CryoBrew (Miltenyi Biotec) + 10% DMSO. PB plasma was obtained by centrifugation and stored at −80°C.

### CyTOF Staining and Acquisition

Cryopreserved MNCs were thawed and diluted dropwise with 25mL of IMDM (ThermoFisher) containing 20% FBS and 0.1mg/mL DNaseI (Scimar). After washing, 1.5 × 10^6 cells were stained for live/dead discrimination using Cell-ID Cisplatin (Fluidigm) in RPMI-1640 for three minutes at room temperature then quenched with FACS buffer (PBS/1% FBS/1mM EDTA). Up to 7 samples, including one of two healthy donor PB MNC samples, were simultaneously barcoded with distinct anti-CD45 metal conjugates and incubated on ice for 30 minutes. Washed cells were then pooled to reduce inter-sample variability. Surface antibody mastermix (Supplemental Table 2, 3) was added to cells and incubated on ice for 30 minutes. Washed cells were fixed with 1 mL of Fixation buffer (Foxp3 Fixation Buffer Kit, eBioscience) for 45 minutes at RT in the dark. Intracellular staining was performed in Permeabilization buffer (Foxp3 Fixation Buffer Kit, eBioscience) with intracellular antibody mastermix (Supplemental Table 2, 3) for 30 minutes at RT, followed by two washes. Cells were resuspended in 200 μL of a 4% paraformaldehyde solution containing 0.125 μM Cell-ID Iridium-191/193 (Fluidigm) and incubated at 4°C overnight. Samples were washed four times with Cell Acquisition Solution (CAS) (Fluidigm), counted, and resuspended in CAS. Data was acquired on a Helios 2 CyTOF instrument at Sydney Cytometry, normalized with EQ Four Element Calibration Beads (Fluidigm), and exported in FCS 3.0 format.

### CyTOF analysis

FCS files were imported into FlowJo Version 10.6 and manually gated to remove normalization beads, doublets and dead cells (cisplatin positive cells), and select live cells (DNA intercalator-bright cells). Individual samples were selected by gating on cells specific for only a single CD45-barcoding antibody and exported from FlowJo as a channel values CSV file. Data were imported into R (Version 4.0.5) and analyzed using the R package ‘Spectre’(*74*). Data were transformed using do.asinh (cofactor = 5), followed by coarse batch correction with run.align (method: ‘95p’). Fine correction was performed using train.cytonorm (goal = ‘mean’; nQ = 101), with the trained model applied via run.cytonorm. Data were clustered using run.flowsom (meta.k = 40). Lastly, data were visualized using the function run.umap with the default parameters. Clusters of interest were isolated and reclustered (meta.k = 20 or 40), excluding cell state markers where possible, and visualized on a UMAP plot. Cluster cell counts per sample were extracted using the table function. A generalized linear mixed model was used to examine the longitudinal changes in cell frequency (SAS version 9.4).

### Single cell sequencing of CD3^+^ cells

Cryopreserved cells were thawed then resuspended in FACS buffer containing DAPI (142 ng/mL). Live cells (DAPI-negative) were sorted on a FACSAria III (BD Biosciences) using a 100 μm nozzle. Sorted cells were counted, pooled to enable SNP-based demultiplexing of patient samples, filtered, and captured on a Chromium Next GEM Single Cell 5’ GEM v2.0. Samples were sequenced on a Illumina Novaseq All samples were previously SNP genotyped to enable demultiplexing, and batch pooling was performed as described (*31*).

### Data Processing, Integration, and Annotation

Single-cell gene expression and TCR V(D)J libraries were processed jointly using Cell Ranger multi (v7.1.0, 10x Genomics), with gene expression reads aligned to GRCh38-2020-A and V(D)J contigs assembled and annotated against the vdj_GRCh38_alts_ensembl-7.1.0 reference. After SNP-based sample demultiplexing, individual datasets were integrated, and batch effects were mitigated using the Harmony algorithm (*75*). Unsupervised clustering and marker gene identification (FindAllMarkers) was performed using Seurat(*76*). Clusters were annotated based on the expression of canonical marker genes, including *CCR7* and *LEF1* (naïve T cells), *SELL* (central memory T cells), *FOXP3* and *IL2RA* (Tregs), *CD4* and *GZMB* (cytotoxic CD4 T cells), *KLRB1* and *TRAV1-2* (MAIT cells), *GZMK*, *IL7R* and *ITGB1* (effector memory CD8 T cells), *CD69*, *S1PR1*^low^ and *KLF2*^low^ (tissue-resident memory T cells), *GZMB* and *ZNF683* (TEMRA), *FCGR3A* and *KLRF1* (NK-like TEMRA), and *TRDV2* (γδ T cells).

### Analysis of T cell single cell data

TCR clonotype analysis and repertoire profiling were conducted using the R package ‘scRepertoire’ (*77*). Clonotypes were defined by VDJ gene usage and the CDR3 nucleotide sequences, enabling the tracking of clonal expansion and diversity across treatment conditions. To characterize TCR specificity, VDJ sequences were aligned against the VDJdb (*34*) database and IEDB (*35*).

Differentially expressed genes (DEGs) were identified using a Logistic Regression framework within the Seurat FindMarkers function. For paired samples, the patient identifier was included as a latent variable to account for inter-patient variability. Pathway enrichment analysis was performed using the MSigDB Hallmark gene set collection via GSEApy (*78*) in “preranked” mode, with genes ranked by log fold change.

TE expression was quantified using scTE (*79*). The RepeatMasker database was used to restrict analysis to well-characterised TE sequences. Aggregate expression scores for TE families (LINE, SINE and LTR) were calculated using the AddModuleScore function within the Seurat R package. Statistical significance for comparisons of continuous variables between pre-treatment and post-treatment groups was determined using a two-tailed paired t-test. For single-cell differential expression, p-values were adjusted for multiple testing using the Bonferroni correction. A p-value < 0.05 was considered statistically significant.

### Analysis of IFN expression in CD34+ cells

Processed gene expression data from bulk CD34^+^ BM cells collected from MDS patients treated with AZA were downloaded from GEO (GSE274999) (*11*). Data was imported into R (v4.2.1), CPM-normalized using edgeR, and then log2-transformed. Data was visualized using ggplot2 (v4.0.2).

Single-cell gene expression data from (*31*) was analysed using Scanpy (v1.10.4). Cells expressing fewer than 200 genes or where mitochondrial reads exceeded 5% of total reads, and genes detected in fewer than 3 cells were removed. Type I, II and III IFN expression was visualized as a violin plot using matplotlib (v3.10.8).

### NULISA data acquisition and analysis

Techniques used for the NULISA assay have been described previously (*38*). Briefly, sample– and target-specific barcodes were quantified and normalized by dividing the target counts + 1 for each sample well by that well’s internal control counts + 1. Inter-plate normalization was conducted by dividing the counts by the medians of the three target-specific inter plate controls and then rescaled by the factor 10^4^. Normalized counts were then log2-transformed to generate NULISA Protein Quantification (NPQ) units. NPQ units from BM samples were further normalised to protein concentration, determined by BCA assay (Pierce). Linear models were fitted using the R (v.4.2.1) package limma (v3.62.2), and moderated t-statistics were computed via empirical Bayes shrinkage of standard errors. All statistical tests were two-sided.

To identify soluble receptors and secreted antagonists that may inhibit cytokine signaling by sequestering their cognate ligands, Claude AI (Anthropic) was used to generate a list of ligand–inhibitor pairs. This list was manually inspected to remove any combinations not supported by the literature. Soluble IL-6R, IL-7R, and IL-15RA were excluded, as these can form circulating complexes that potentiate rather than inhibit signaling (*80–82*). As NPQ values are reported on a log₂ scale, subtraction of the inhibitor NPQ from the cytokine NPQ yields a log₂ ratio of cytokine to inhibitor abundance; a negative net value indicates that inhibitor abundance exceeds ligand abundance, suggestive of active sequestration. Ratio values were used as input for differential abundance analysis performed using the R package limma.

Spearman correlations between the change in growth factor abundance and the change in blast abundance or cytotoxic CD8 T cell abundance were calculated using scipy.stats.spearmanr (v1.16.3) in Python (v3.12.4). Correlations above 0.5 were visualized on a heatmap made using the Seaborn package (v0.13.2).

### BM trephine imaging and analysis

Trephine biopsies were fixed and decalcified using one of three protocols: formalin-acetic acid-alcohol (FAA) fixation for 3 hours followed by trichloroacetic acid decalcification for 90 minutes, FAA-fixation for 3 hours followed by EDTA decalcification for 24 hours, or Bouin’s fixative followed by decalcification in 10% EDTA in 0.1M Tris-HCl pH 7.

Formalin-fixed paraffin-embedded tissue sections were stained using an 8-plex Opal multiplex immunofluorescence protocol on the Leica Bond RX platform (Leica). Slides were dewaxed using Bond Dewax Solution (Leica) followed by heat-induced epitope retrieval (ER2, 100°C, 20 min). Endogenous peroxidase activity was quenched with peroxide prior to antibody staining. Slides were incubated with a blocking solution comprising Antibody Diluent/Blocking Buffer (AKOYA), 10% goat serum, and 1:100 Human TruStain FcX, (BioLegend). Sequential rounds of staining were performed using primary antibodies against Ki67, PD-1, CD56, CXCL12, CD4, CD3, CD34, CD8, with antibody-specific antigen retrieval conditions and incubation times (Supplemental Table 6). Signal amplification was achieved using Opal fluorophores (Opal 540, 620, 570, 690, 480, 520, 650, and 780; Akoya Biosciences) with antibody stripping using ER1 or ER2 retrieval steps (95°C, 20 min) between staining cycles. Detection was performed using Novolink polymer reagents (Leica), and nuclei were counterstained with spectral DAPI prior to imaging on AKOYA Phenoimager (AKOYA). Images were acquired as overlapping tiles.

Image tiles were stitched together and analyzed using Halo AI (Indica Labs, v3.6.4). A custom nuclear classifier was trained on manually annotated nuclei and background regions from 4 representative slides which were manually chosen (resolution: 0.25 µm/px; minimum object size: 5 µm). Detected nuclei were used to define individual cells, and a cell phenotype classifier was subsequently trained using the object phenotyper module. As the object phenotyper incorporates tissue context into cell classification, training data included both isolated and densely clustered cells from multiple trephines to ensure accurate generalization across samples. All cell locations and identities were exported from Halo AI as tables.

### Neighborhood analysis

For each focal cell, a local neighborhood composition matrix was generated to quantify the spatial composition of surrounding cell types within a 100 µm radius. The focal cell was excluded from this calculation. For every neighboring cell, the Euclidean distance to the focal cell was calculated and converted to a proximity-weighted contribution using an inverse exponential weighting function:

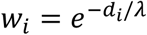

where *d_i_* is the Euclidean distance from the focal cell to neighbouring cell *i*, and *λ* was set to one-third of the search radius (*λ* = 33.3µm for a 100 µm radius). Weighted contributions were summed separately for each unique neighbouring cell type:

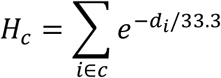

where *H_c_* is the weighted neighborhood *over all neighbouring cells of type c*. This produces a continuous local composition profile in which nearby cells contribute more strongly than cells further from the focal cell, rather than using raw neighbour counts.

Neighborhood composition features were standardized using z-score normalization implemented in Scikit-learn (1.7.2) prior to dimensionality reduction. Uniform Manifold Approximation and Projection (UMAP) was then applied to the standardized feature matrix using Euclidean distance as the similarity metric with 15 nearest neighbours, a minimum distance parameter of 0.1, and a fixed random seed of 42 using UMAP-learn (0.5.9.post2). The resulting two-dimensional embedding was used for visualization and downstream clustering. To identify neighborhood states, a k-nearest neighbour graph was constructed from the UMAP coordinates using 100 nearest neighbours per cell with the kneighbors_graph implementation in Scikit-learn (1.7.2). Leiden community detection was subsequently performed on this graph using IGraph (0.11.9) and Leidenalg (0.10.2), employing the RBConfigurationVertexPartition implementation with a resolution parameter of 0.5. Clustering was iterated until convergence and initialized with a random seed of 42 to ensure reproducibility. Cluster assignments were subsequently mapped back to the original dataset and visualized on the UMAP embedding. The approximate boundary of the UMAP embedding (used as a visualization aid in Fig.6 D-J) was generated as a kernel density estimate using a contour at 5% of the maximum estimated density and manually refined.

Statistical analysis of neighborhood frequency in responders and non-responders was performed using a generalized-linear mixed model in SAS v 9.4. Bar heights throughout show the mean, error bars show standard deviation, and box plots show median and interquartile range (plus minimum and maximum values (excluding outliers) in Fig. 3 and S Fig. 3). Custom code is available at https://github.com/henryrhampton/Bone-marrow-microenvironment-in-MDS

### List of Supplemental Materials

Supplemental Figure 1. GMPs expanded when patients responded to AZA.

Supplemental Figure 2. Cytotoxic CD8 T cells expanded when patients responded to AZA.

Supplemental Figure 3. Limited expression of IFN genes in patient CD8 T cells and CD34^+^ HSPCs.

Supplemental Figure 4. CD8 T cells showed limited reactivity to known cancer-associated antigens.

Supplemental Figure 5. Robust detection of molecules in blood and bone marrow using NULISA technology.

Supplemental Figure 6. Ki67 and PD-1 expression in HSPCs and T cells in bone marrow trephine sections.

Supplemental Figure 7. Neighborhood composition differs between responders and non-responders at clinical response.

Supplemental Table 1. Patient characteristics and inclusion in analysis modalities.

Supplemental Table 2. Myeloid-focused CyTOF panel.

Supplemental Table 3. T cell-focused CyTOF panel.

Supplemental Table 4. Plasma changes.

Supplemental Table 5. Features of neighborhood clusters.

Supplemental Table 6 Antibodies for Opal9 immunohistochemistry.

## Acknowledgments

We thank Eric Lam, Dr Winnie Lau, and Dr Alejandro Rios Villamil for their contributions to the single-cell experiments, Jessica Root for assistance with the NULISA assay, Dr Chan Phetsouphanh (UNSW Sydney), and Dr John Zaunders (St Vincent’s Hospital, Sydney), A/Prof Sunil Kumar Saini (DTU, Copenhagen), Prof. Austin Kulasekararaj (KCL, UK), Prof. Fabio Luciani (UNSW Sydney), and A/Prof. Fabio Zanini (UNSW Sydney) for insightful discussions. Biospecimens and data used in this research were obtained from the Health Precincts Biobank, UNSW Biospecimen Services, UNSW Sydney, Australia. The Biobank acknowledges the key contribution of NSW Health Pathology.

## Funding

National Health and Medical Research Council, grant GNT2011627 to J.E.P, C.J.J and H.R.H

Anthony Rothe Memorial Trust to J.A.I.T.

Anthony Rothe Memorial Trust to J.E.P.

Medical Research Future Fund, grant MRF1200271 to J.E.P, M.N.P and J.O.

Cancer Institute of New South Wales, grant 2022/TPG2152 to J.E.P, M.N.P, J.O and J.A.I.T

## Author contributions

Conceptualization: J.E.P.

Investigation: H.R.H., A.P, M.C., B.W., D.S., M.K., I.S., S.J., M.N.T.N, F.Y., S.D., N.C., N.T., J.O.

Funding acquisition: H.R.H., M.N.P, J.O., J.A.I.T, C.J.J, J.E.P.

Supervision: J.W.H.W, M.N.P, H.M.G, H.A.A, A.J., J.O., J.A.I.T, C.J.J, J.E.P.

Writing – original draft: H.R.H., J.A.I.T, J.E.P. Essential reagents: D.K.H., M.T.

Writing – review & editing: H.R.H., J.A.I.T, C.J.J, J.E.P.

## Supplemental Figures and Legends

**Supplemental Figure 1.**
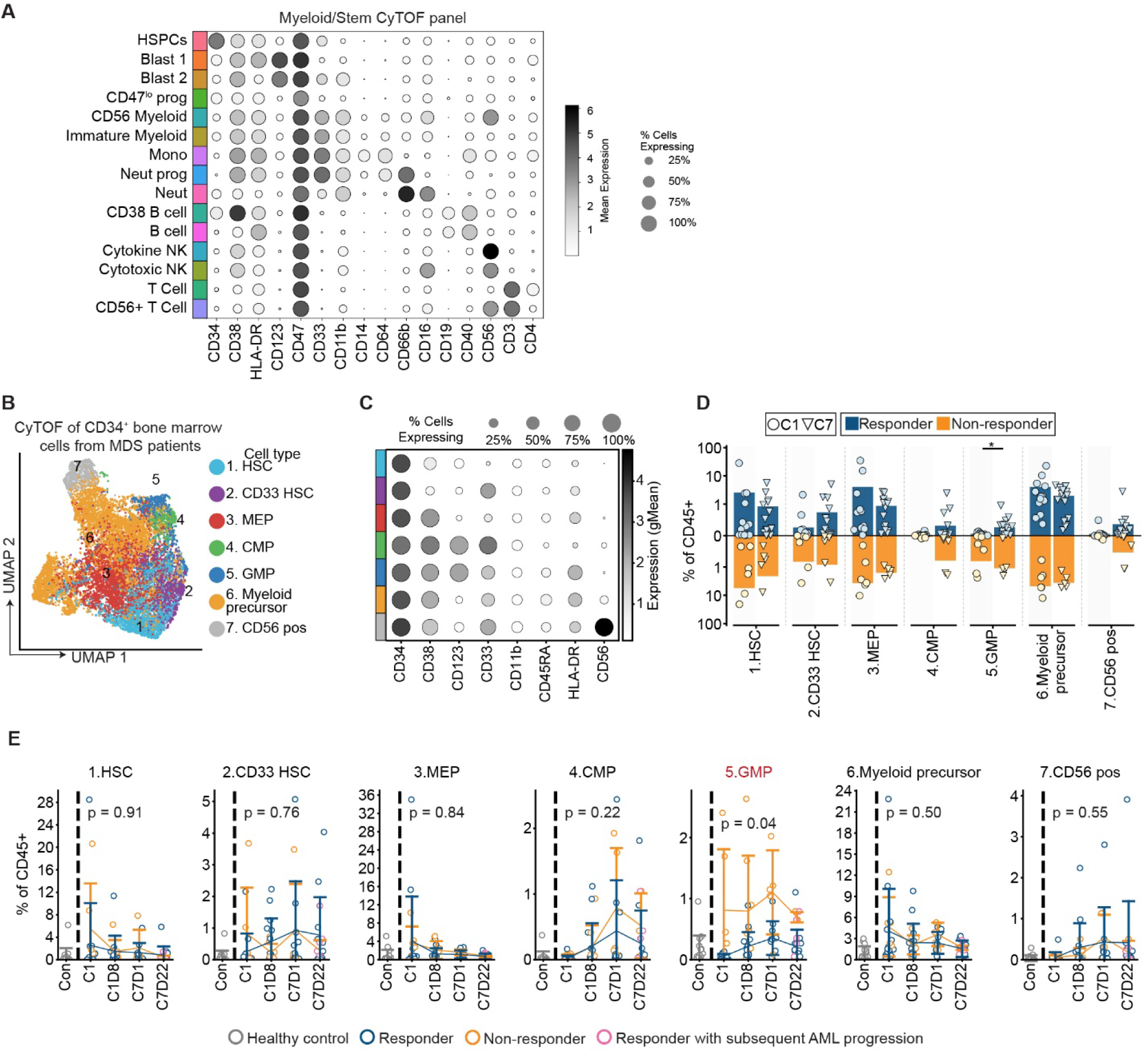
GMPs expanded when patients responded to AZA. (**A**) Abundance of surface proteins used to annotate cell types detected in the Myeloid/Stem CyTOF panel. (**B**) UMAP embedding showing CD34^+^ subsets identified by mass cytometry. (**C**) Abundance of selected surface proteins on CD34^+^ subsets. (**D**) Changes in CD34^+^ subset frequencies between diagnosis (C1) and C7. P-values: linear mixed-effects model, bar height shows the mean. (**E**) Longitudinal changes in abundance of specified HSPC subsets. Pink markers indicate responders with subsequent disease progression. P-values: generalized linear mixed model. Error bars show standard deviation. *** p < 0.001.

**Supplemental Figure 2.**
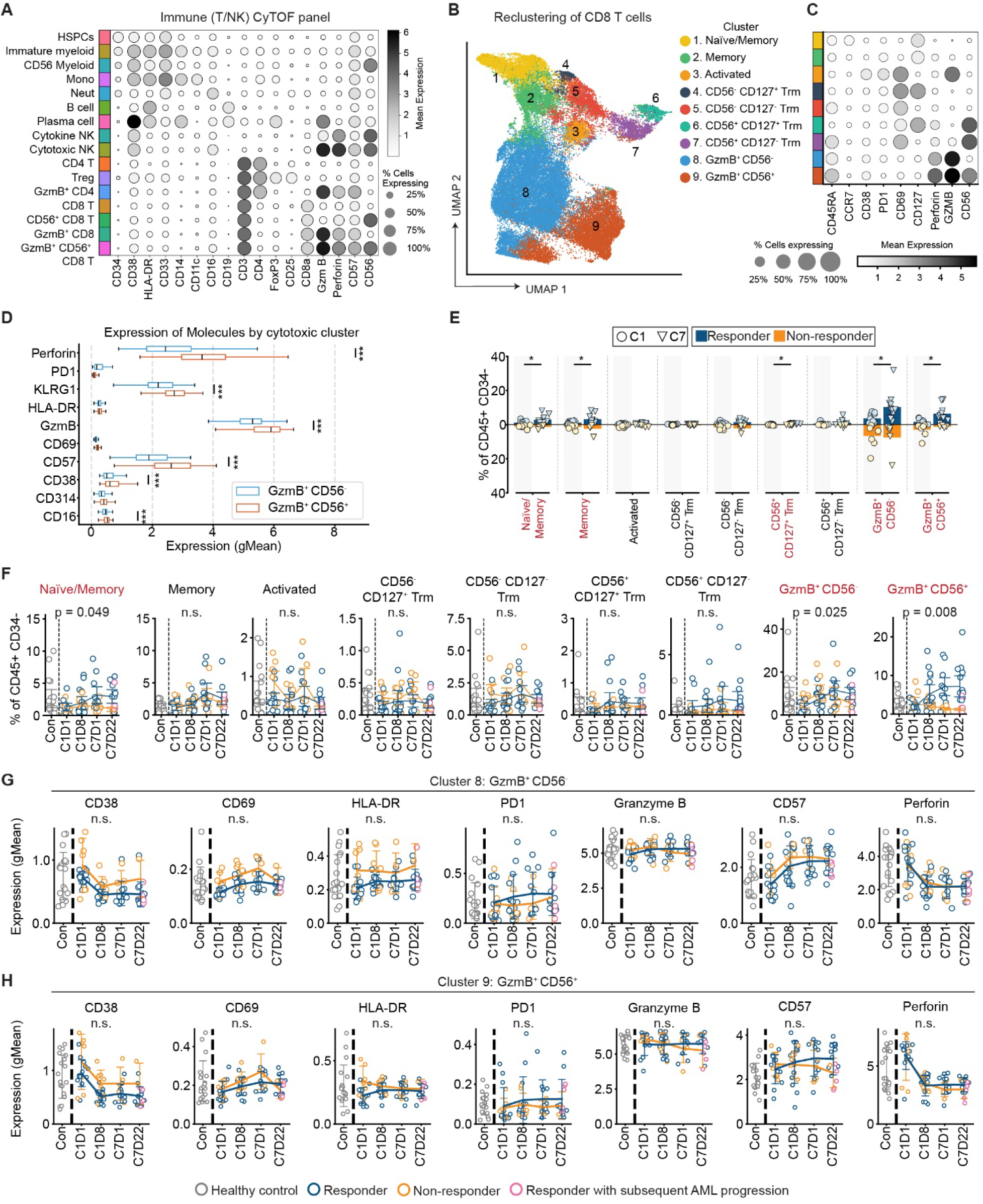
Cytotoxic CD8 T cells expanded when patients responded to AZA. (**A**) Abundance of proteins used to annotate cell types detected in the Immune (T/NK) CyTOF panel. (**B**) UMAP embedding showing CD8 T cell clusters identified by FlowSOM. (**C**) Abundance of selected surface proteins on CD8 T cell clusters. (**D**) Expression of selected molecules on CD56^-^GzmB^+^CD8 T cells and CD56^+^GzmB^+^CD8 T cells. P-values: paired t-test. (**E**) Changes in CD8 T cell cluster abundance between diagnosis and C7. P-values: linear mixed-effects model, bar height shows the mean. (**F-H**) Longitudinal changes in abundance of (**F**) CD8 T cell clusters, (**G-H**) molecules associated with T cell activation and cytotoxicity in (**G**) CD56^-^GzmB^+^CD8 T cells, and (**H**) CD56^+^GzmB^+^CD8 T cells. Pink markers indicate responders with subsequent disease progression. P-values: generalized linear mixed model. Error bars show standard deviation. * p < 0.05, *** p < 0.001.

**Supplemental Figure 3.**
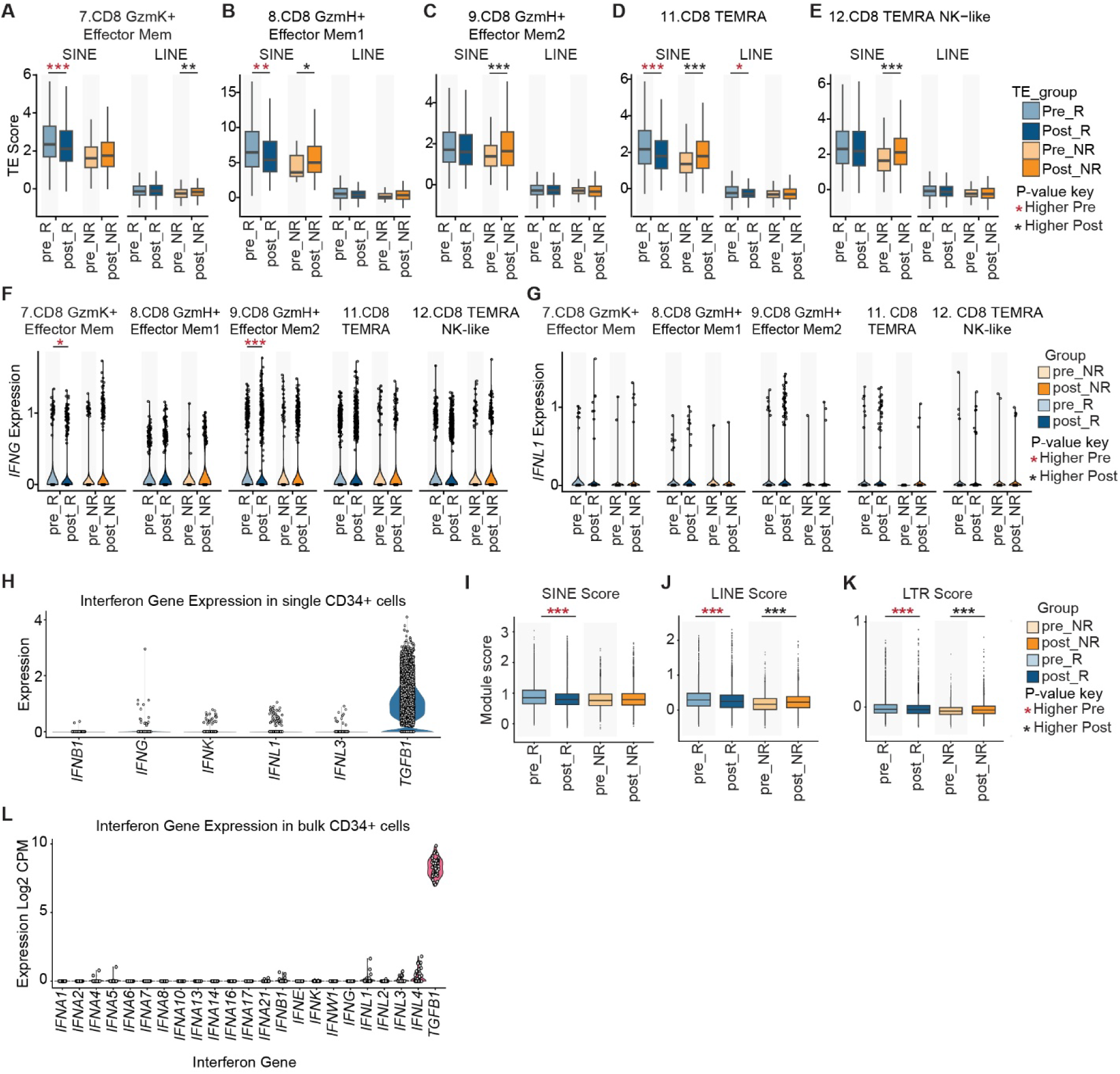
Limited expression of IFN genes in patient CD8 T cells and CD34^+^ HSPCs. (**A-E**) Expression of SINE and LINE elements in selected CD8 T cell clusters. P value: Mann–Whitney U test with Benjamini–Hochberg correction for multiple comparisons. (**F-G**) Expression of *IFNG* (**F**) and *IFNL1* (**G**) in selected CD8 T cell clusters. P value: Mann–Whitney U test with Benjamini–Hochberg correction for multiple comparisons. (**H**) Expression of IFN genes (plus positive control gene *TGFB1*) in *34,494* single CD34^+^ cells from patients with HR-MDS (*31*). (**I-K**) Expression of (**I**) SINE, (**J**) LINE, and (**K**) LTR elements in 34,494 single CD34^+^ cells from patients with HR-MDS (*31*). P value: Mann–Whitney U test with Benjamini–Hochberg correction for multiple comparisons. (**L**) Expression of IFN genes and *TGFB1* by bulk sequencing in magnetically isolated CD34^+^ cells from HR-MDS patients (*11*), with 52 samples collected across multiple timepoints. * p <= 0.05, ** p <= 0.01, *** p <= 0.001. Color of asterisks indicates the direction of change from pre– to post-treatment.

**Supplemental Figure 4.**
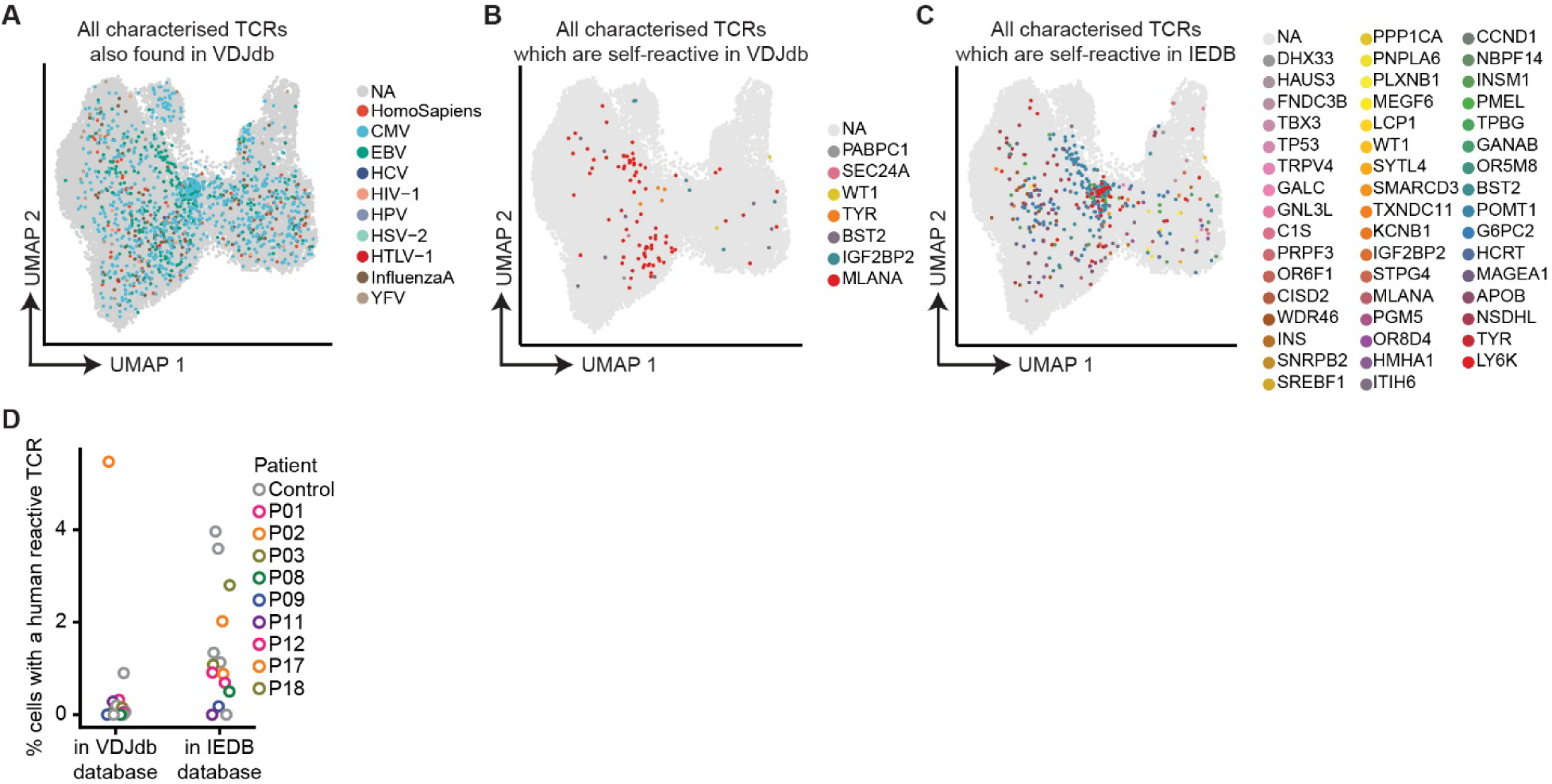
CD8 T cells showed limited reactivity to known cancer-associated antigens. (**A**) UMAP embedding colored by the presence of TCR sequences with known antigen reactivity as annotated in VDJdb (*83*). (**B-C**) UMAP embedding colored by TCR reactivity for self– or cancer-associated antigens, annotated using (**B**) VDJdb (*83*) or (**C**) IEDB (*35*). (**D**) Percent of CD8 T cells with a self-reactive TCR in VDJdb and IEDB.

**Supplemental Figure 5.**
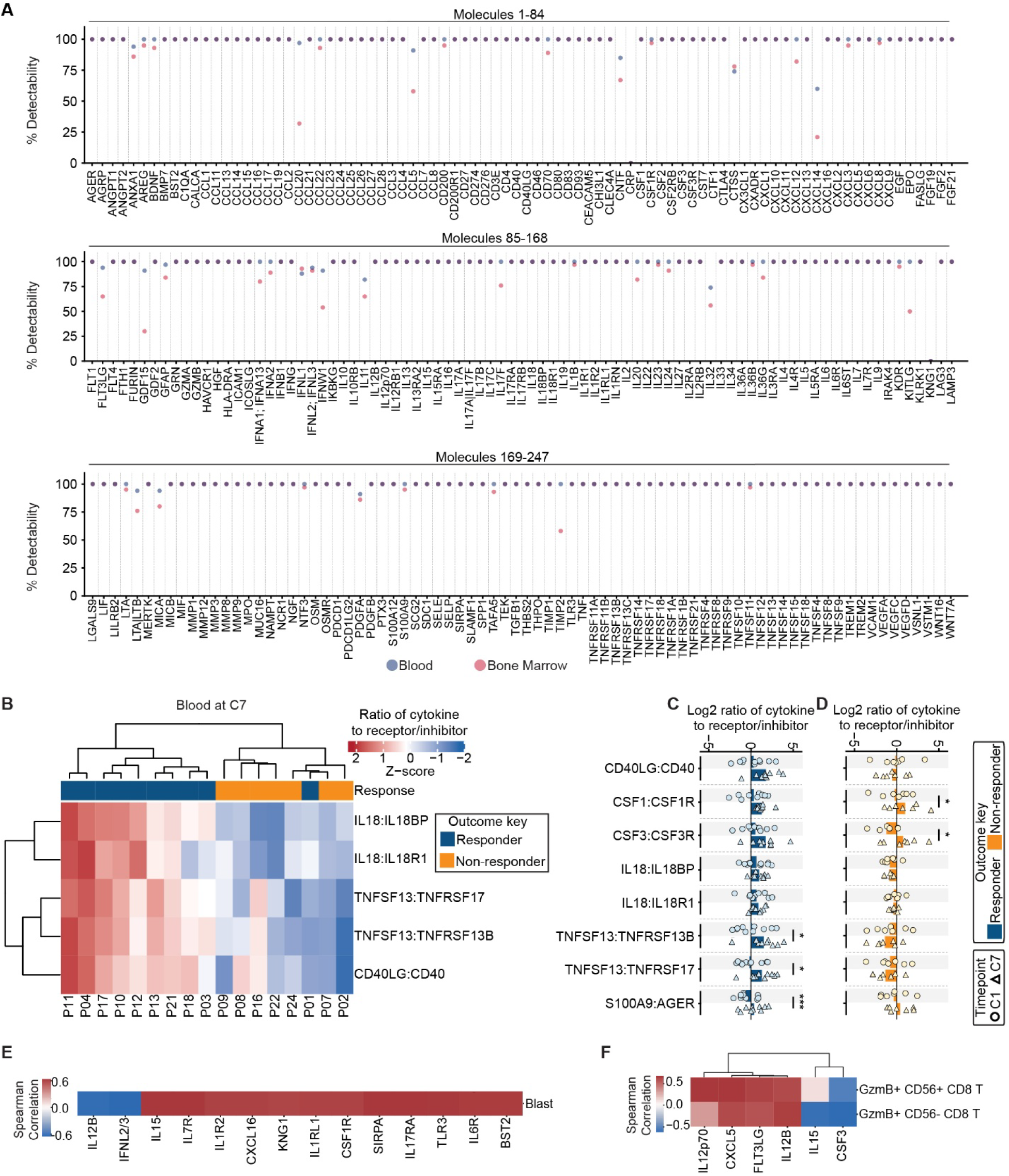
Robust detection of molecules in blood and bone marrow using NULISA technology. (**A**) Percentage of samples where each analyte was detected above background. (**B**) Ratio of selected cytokine to receptor/inhibitor pairs at C7 in blood. (**C-D**) Ratio of selected cytokine to receptor/inhibitor pairs at C1 and C7 in the blood of (**C**) responders, and (**D**) and non-responders. P-values: linear mixed-effects model bar height indicates mean. (**E-F**) Spearman correlation of change in cytokine abundance to change in (**E**) blast percentage, or (**F**) cytotoxic CD8 T cell subset percentage.

**Supplemental Figure 6.**
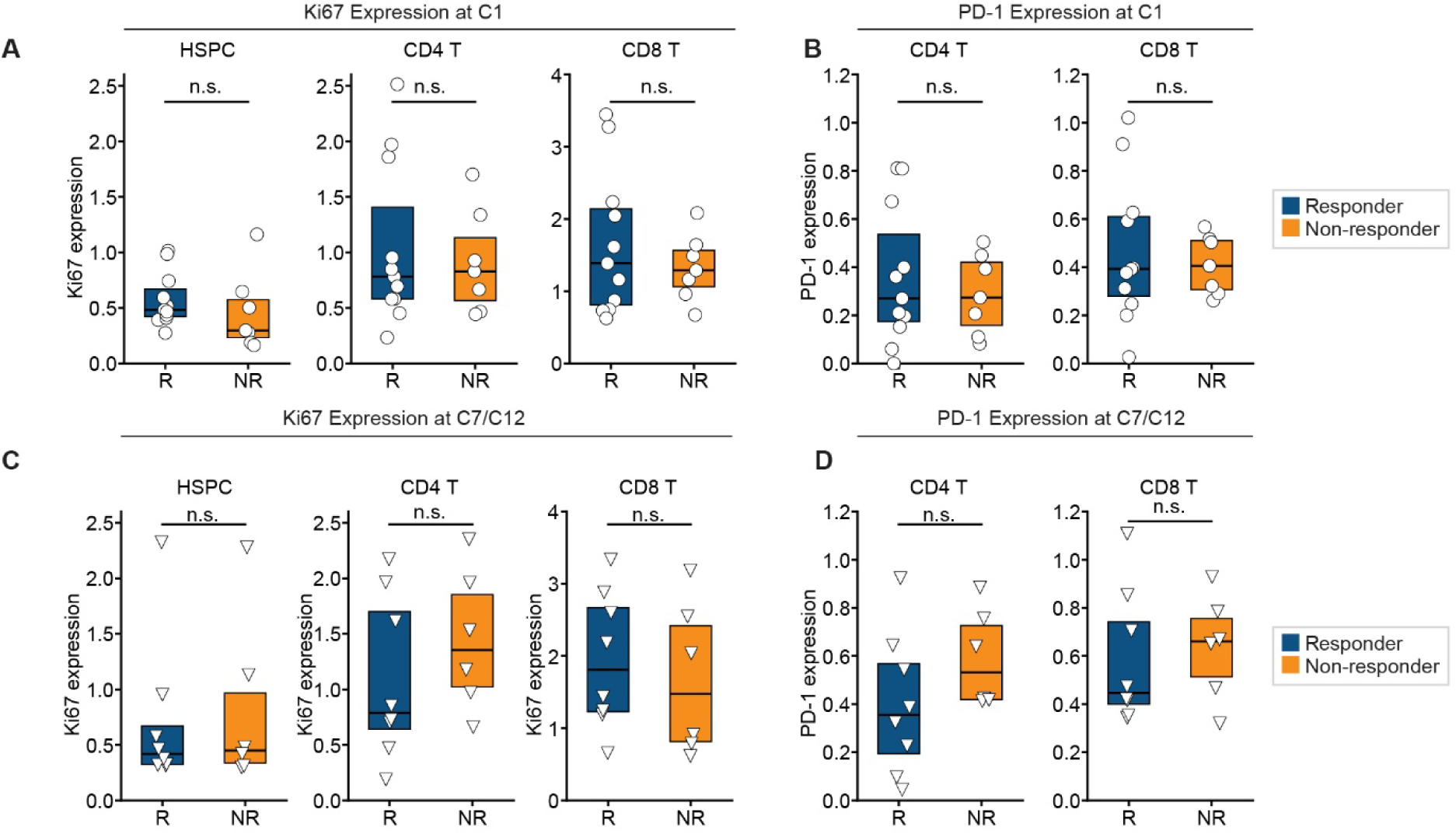
Ki67 and PD-1 expression in HSPCs and T cells in bone marrow trephine sections. (**A**) Ki67 expression in HSPCs and CD4 and CD8 T cells at diagnosis. (**B**) PD-1 expression in CD4 and CD8 T cells at diagnosis. (**C**) Ki67 expression in HSPCs and CD4 and CD8 T cells at C7/C12. (**D**) PD-1 expression in CD4 and CD8 T cells at C7/C12. P-values: Mann-Whitney U test.

**Supplemental Figure 7.**
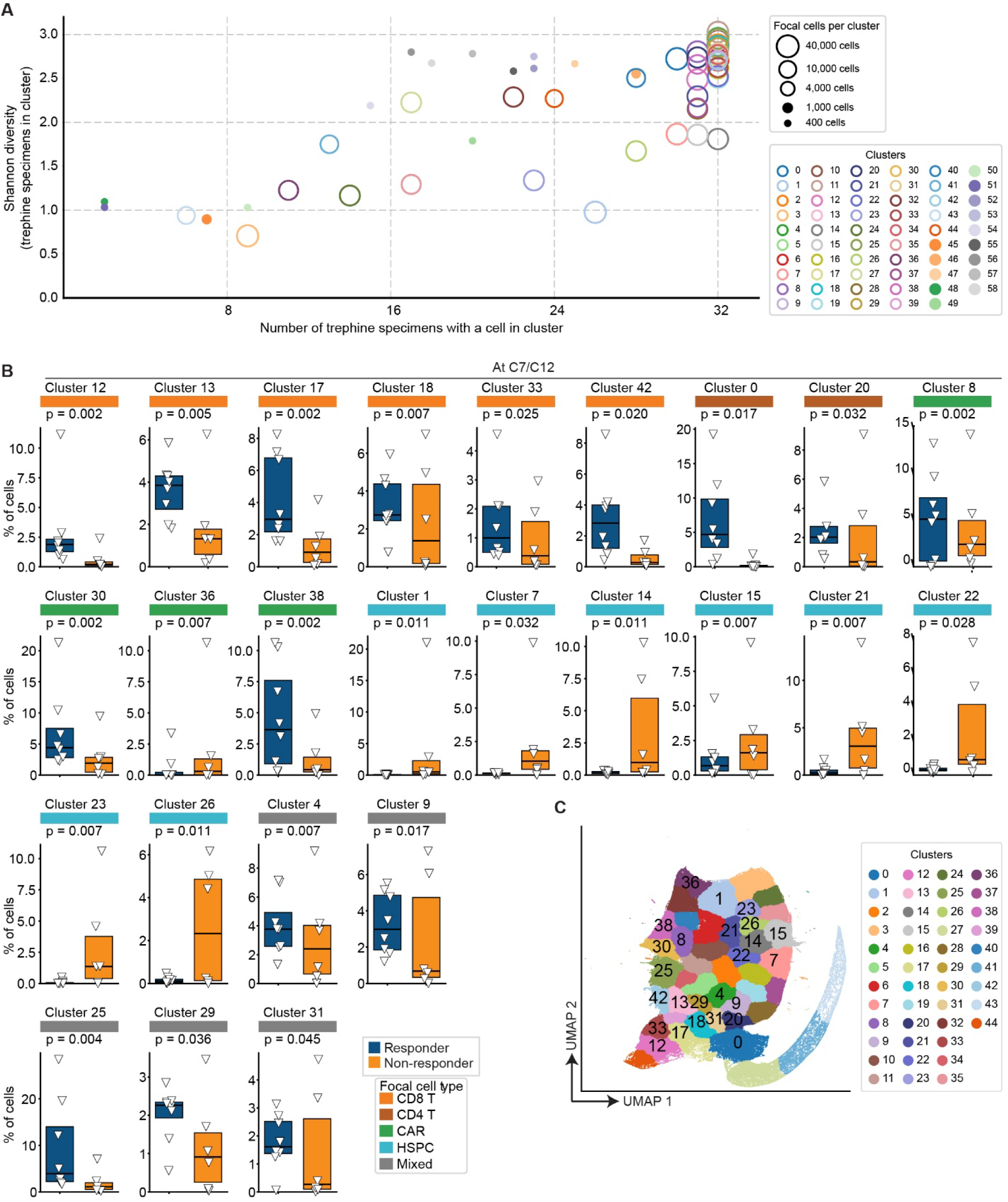
Neighborhood composition differs between responders and non-responders at clinical response. (**A**) Number of contributing trephines (x axis), Shannon diversity of contributing trephines (y axis), and number of focal cells (circle size) for each proximity-weighted neighborhood cluster. Clusters containing less than 1000 cells (filled circles) were excluded from further analysis. Higher Shannon diversity indicates that neighborhoods within the cluster are drawn relatively evenly from different trephine specimens, while lower values indicate dominance by one or a few specimens. (**B**) Percentage of cells from each trephine specimen found in specific neighborhoods at clinical response (C7/C12). P-values: generalized linear mixed model. (**C**) UMAP embedding of focal cells colored by proximity-weighted neighborhood cluster and labelled to locate clusters listed in **B**.

